# Hypothalamic circuitry underlying stress-induced insomnia and peripheral immunosuppression

**DOI:** 10.1101/2020.04.29.069393

**Authors:** Shi-Bin Li, Jeremy C Borniger, Hiroshi Yamaguchi, Julien Hédou, Brice Gaudillière, Luis de Lecea

## Abstract

The neural substrates of insomnia/hyperarousal induced by stress remain unknown. Here, we show that restraint stress leads to hyperarousal associated with strong activation of corticotropin-releasing hormone neurons in the paraventricular nucleus of hypothalamus (CRH^PVN^) and hypocretin neurons in the lateral hypothalamus (Hcrt^LH^). CRH^PVN^ neurons are quiescent during natural sleep-wake transitions but are strongly active under restraint stress. CRISPR-Cas9-mediated knockdown of the *crh* gene in CRH^PVN^ neurons abolishes hyperarousal elicited by stimulating LH-projecting CRH^PVN^ neurons. Genetic ablation of Hcrt neurons or *crh* gene knockdown significantly reduces insomnia/hyperarousal induced by restraint stress. Given the association between stress and immune function, we used single-cell mass cytometry by time of flight (CyTOF) to analyze peripheral blood and found extensive changes to immune cell distribution and functional responses during wakefulness upon optogenetic stimulation of CRH^PVN^ neurons. Our findings suggest both central and peripheral systems are synergistically engaged in the response to stress via CRH^PVN^ circuitry.

## INTRODUCTION

A brief exposure to a stressor imposes a persistent imprint on the brain (*1*). Insomnia is among the most prevalent stress-related complaints, and clarification of the mechanism underlying stress-induced insomnia is essential for developing effective treatment. Accumulated evidence has shown that wakefulness-promoting brain regions are involved in the neural response to stress (*2*). Additionally, changes in adaptive and maladaptive immune responses has been empirically linked to stress exposure (*3*). However, the explicit neural circuitry integrating behavioral (e.g., insomnia) and physiological (e.g., immunosuppression) effects of stress remains unclear.

Stress has been known to induce insomnia/hyperarousal in humans (*4, 5*) and rodent models (*2*). Transitions from sleep to wakefulness can be elicited upon optogenetic activation of the hypocretin/orexin (Hcrt) system (*6*), noradrenergic locus coeruleus (LC) (*7, 8*), dopaminergic neurons in ventral tegmental area (VTA) (*9*) and dorsal raphe (*10*), and/or the cholinergic basal forebrain (BF), among others (*11, 12*). Despite the strong evidence demonstrating a causal relationship between neuronal ensembles and behavioral wakefulness, whether these same cell populations are recruited during stress exposure remains unknown. Indeed, some arousal-promoting brain nuclei are preferably active during appetitive sensory inputs. For example, fiber photometry recording from dopaminergic neurons in the VTA (*9*) and dorsal raphe nucleus (*10*) in male mice showed maximum activity during presence of palatable food and female mice suggesting their association to positive valence. Furthermore, the observation that reward buffers the CRH^PVN^ neuronal activities following footshock stressor suggests that the reward-related neuronal ensembles are unlikely implicated in stress circuitry (*13*). Interestingly, both positive and negative stimuli are able to elicit Hcrt neuronal activities (*14*). These studies indicate different arousal-promoting brain nuclei may mediate arousal to various stimuli depending on their respective valence (either positive or negative). A well-defined neural circuit linking stress to arousal has yet to be adequately described.

Psychosocial stress has well known effects on systemic immune responses (*3, 15*). Specifically, chronic stressors suppress both cellular and humoral measures (*16*). For example, in a classic study of chronically stressed dementia patient caregivers, Kiecolt-Glaser and colleagues observed dramatic decrements in three separate measures of cellular immunity (*17*). In animal models, psychological stress (e.g., via restraint stress) causes thymic involution (along with T-cell apoptosis) and suppresses granulocyte and macrophage migration to the location of inflammatory challenge, putatively via a glucocorticoid-dependent mechanism (*18, 19*). Indeed, a major driver of immune responses to stressors are glucocorticoids secreted from the adrenal cortex, downstream of the hypothalamic-pituitary-adrenal (HPA) axis (*20*). CRH^PVN^ neurons serve as the initial node in this axis, transducing the neuronal signal of stress into an endocrine output (i.e., glucocorticoid secretion). The actions of the HPA axis are fundamentally coupled to arousal, as sleep disruption powerfully drives HPA activation and glucocorticoid secretion in human (*21*). Despite the clear effects of stress and stress-induced arousal on the immune system, an understanding of the neural circuitry underlying this phenomenon is lacking.

We hypothesized that insomnia/hyperarousal and peripheral immunosuppression may share the same neural substrates under stress. We performed a series experiments with transgenic mice to test this hypothesis. After identifying that both stress-associated CRH^PVN^ neurons and arousal-promoting Hcrt^LH^ neurons are activated by restraint stress, we demonstrated that Hcrt^LH^ neurons are monosynaptically innervated by CRH^PVN^ neurons. We then observed that the activity of CRH^PVN^ neurons is not required for natural sleep-to-wake transitions, but associates with stress exposure. Mild optogenetic stimulation of CRH^PVN^, but not Hcrt^LH^ neurons, elicited persistent wakefulness mimicking insomnia induced by restraint stress. Interestingly, the absence of Hcrt neurons or the *crh* gene in CRH^PVN^ neurons counteracted insomnia/hyperarousal induced by optogenetic stimulation or restraint stress. Unbiased CyTOF analysis revealed optogenetic stimulation of CRH^PVN^ neurons leads to peripheral immunosuppression simulating the physiological effect due to stress. These data suggest that connections between CRH^PVN^ and Hcrt^LH^ neurons drives stress-induced insomnia, and that activation of CRH^PVN^ neurons is sufficient to elicit stress-induced immunosuppression.

## RESULTS

### Restraint stress causes strong cFos expression in CRH^PVN^ and Hcrt^LH^ neurons

To reveal the specific neuronal responses to stress exposure, we investigated the stress-related paraventricular nucleus of the hypothalamus (PVN) and the arousal-related brain region the lateral hypothalamus (LH). We subjected mice to restraint stress, a well-established robust and non-invasive stress paradigm (*22*), and performed cFos antibody staining at different time points after restraint stress. We found a large number of cells around the PVN and LH to be cFos-positive (Fig. S1) consistent with earlier work showing cFos activity upon exposure to cage exchange stress in rats (*2*). Antibody staining in the same slices revealed that the vast majority of PVN cFos-positive neurons co-express corticotropin-releasing hormone (CRH, also known as corticotropin-releasing factor (CRF)) and a major population of LH cFos-positive neurons are hypocretin-1 (Hcrt1)-positive (Fig. S1), a group of neurons that play a major role in promoting and stabilizing wakefulness (*6*).

### CRH^PVN^ neurons directly innervate Hcrt^LH^ neurons

The PVN consists of a constellation of several neuronal types including those that secrete the neuropeptides CRH, AVP, and OXT (*23*). It was unclear which cell type in the PVN directly makes synapses onto Hcrt^LH^ neurons, despite previous reports that the PVN is an upstream partner of Hcrt neurons (*14, 24*). We used the EnvA-pseudotyped glycoprotein (G)-deleted rabies virus (EnvA + RV*d*G) (*25, 26*) to trace the monosynaptic inputs to Hcrt neurons and found a major population of PVN neurons directly project to Hcrt neurons (*14*). We further performed antibody staining in the PVN and found around 60% of the RV*d*G-labeled PVN neurons are CRH-positive and about half of the CRH^PVN^ neurons were labeled by the RV*d*G (Fig. 1A-E). Interestingly, arginine vasopressin (AVP) and oxytocin (OXT) neurons, another two major populations of neurons intermingled with CRH neurons in the PVN (*23*) were labeled with a much lower ratio (Fig. 1F-J & K-O). We then injected AAV-Retro vectors (*27*) carrying eYFP to the LH of CRH::Cre, AVP::Cre and OXT::Cre mouse lines respectively and observed that nearly 70% of CRH neurons project to the LH containing Hcrt neurons (Fig. S2A-E). Of note, also about 60% AVP neurons were labeled by AAV-Retro vectors (Fig. S2F-J) suggesting AVP^PVN^ neurons may target neuronal populations that are distinct from Hcrt neurons in the LH given cell type-specific monosynaptic tracing of Hcrt neurons in Fig. 1. These two sets of experiments demonstrated CRH^PVN^ neurons are the primary population targeting Hcrt^LH^ neurons, and established a direct link between neurons orchestrating stress and arousal. Interestingly, both tracing strategies identified only limited OXT neurons projected to the LH (Fig. 1K-O & Fig. S2K-O) confirming earlier observations (*28*).

**Fig. 1.**
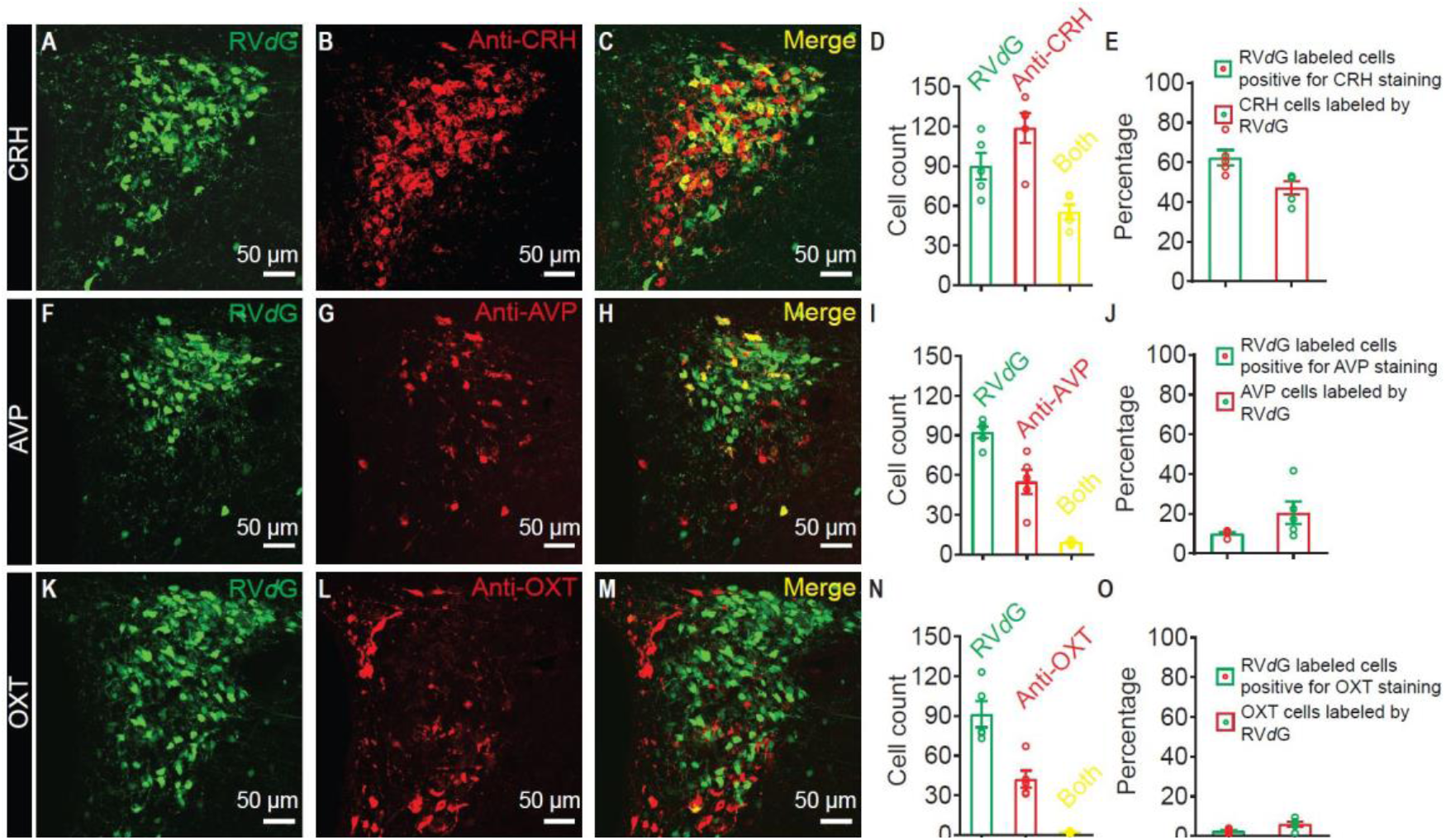
CRH^PVN^ neurons project to Hcrt^LH^ neurons with monosynaptic contacts in Hcrt::Cre mice. (**A**- **E**) Representative slice containing PVN neurons labeled by RV*d*G (**A**), antibody staining against CRH (**B**), merged slice (**C**), cell counts of RV*d*G labeled neurons, CRH-positive neurons and both-positive neurons (**D**), and percentages of RV*d*G labeled neurons positive for CRH staining and CRH neurons labeled by RV*d*G (**E**). (**F-J**), Representative slice containing PVN neurons labeled by RV*d*G (**F**), antibody staining against AVP (**G**), merged slice (**H**), cell counts of RV*d*G labeled neurons, AVP-positive neurons and both-positive neurons (**I**), and percentages of RV*d*G labeled neurons positive for AVP staining and AVP neurons labeled by RV*d*G (**J**). (**K-O**) Representative slice containing PVN neurons labeled by RV*d*G (**K**), antibody staining against OXT (**L**), merged slice (**M**), cell counts of RV*d*G labeled neurons, OXT-positive neurons and both-positive neurons (**N**), and percentages of RV*d*G labeled neurons positive for OXT staining and OXT neurons labeled by RV*d*G (**O**) (n = 5 for each group).

### Activity of CRH^PVN^ neurons is not a prerequisite for natural sleep-to-wake transitions

We then monitored the activity of CRH^PVN^ neurons in CRH::Cre mice and Hcrt^LH^ neurons in Hcrt::Cre mice by measuring Ca^2+^ transients during natural sleep/wake transitions using fiber photometry (Fig. 2). Interestingly, CRH^PVN^ neurons show minimal changes in activity during natural sleep/wake transitions (Fig. 2A-C). However, Hcrt^LH^ neurons are highly active during wakefulness (Fig. 2D-F), confirming earlier findings (*29, 30*). These experiments demonstrate that activation of Hcrt neurons does not require CRH^PVN^ neuronal activity during natural sleep-to-wake transitions.

**Fig. 2.**
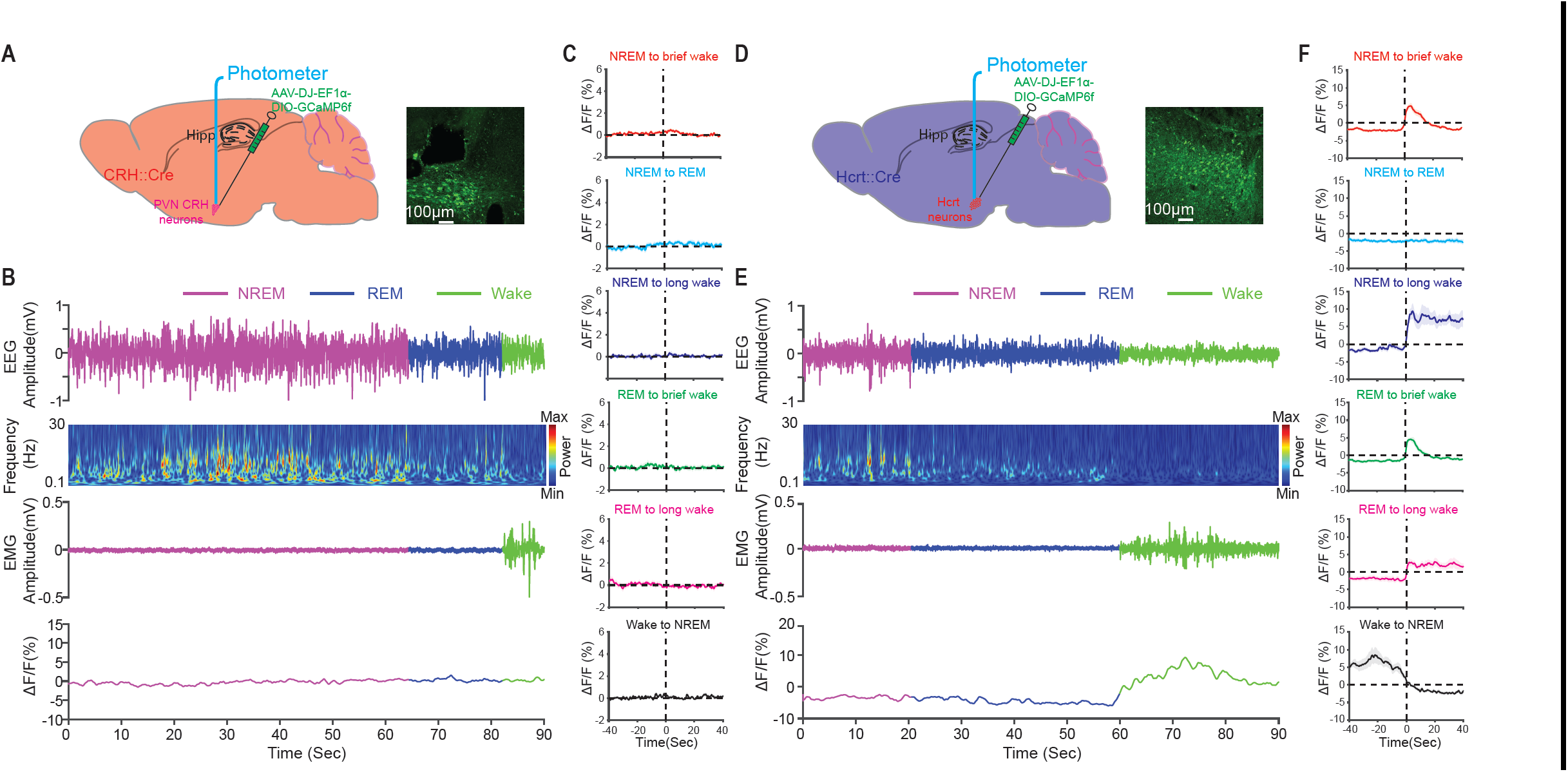
CRH^**PVN**^ **and Hcrt**^**LH**^ **neuronal activity during natural sleep/wake transitions and exposure to salient stimuli**. (**A**) Schematic of fiber photometry recording from CRH^PVN^ neurons. (**B**) Representative EEG, EEG power spectra, EMG and CRH^PVN^ GCaMP trace recorded simultaneously during a 90 sec natural sleep/wake transition episode from a CRH::Cre mouse. (**C**) No obvious fluctuations of Ca^2+^ transients in CRH^PVN^ neurons during natural sleep/wake transitions. (**D**) Schematic of fiber photometry recording from Hcrt^LH^ neurons. (**E**) Representative EEG, EEG power spectra, EMG and Hcrt^LH^ GCaMP trace recorded simultaneously during a 90 sec natural sleep/wake transition episode from a Hcrt::Cre mouse. (**F**) Strong Ca^2+^ transients correlate with wakefulness in Hcrt^LH^ neurons during natural sleep/wake transitions. (n = 7 for **A-C**, n = 6 for **D-F**).

### Stress elicits strong CRH^PVN^ and Hcrt^LH^ neuronal activities

We further investigated how CRH^PVN^ and Hcrt^LH^ neurons change their activity in response to various salient stimuli including novelty, social, appetitive and aversive components. Generally, aversive stimuli such as 2, 3, 5-Trimethyl-3-thiazoline (TMT, a component of fox odor), restraint stress, open arm of the elevated plus maze (EPM), and manual grabbing triggered significant CRH^PVN^ neuronal activity, whereas neutral or appetitive stimuli including novelty (novel Falcon tube), female mice, high fat diet (HFD) elicited CRH^PVN^ neuronal activity to a lesser extent (Fig. S3A&B). The increase of CRH^PVN^ neuronal activity upon aversive stimuli is consistent with previous work using aversive stimuli including forced swim test, tail-restraint test, overhead object, TMT, looming disk (*31*), and white noise (*32*). Interestingly, Hcrt^LH^ neurons displayed strong neuronal activity to various stimuli regardless of stimulus valence (Fig. S3C&D). Of note, restraint stress triggered robust activity in both of CRH^PVN^ and Hcrt^LH^ neurons especially at the beginning of restraint stress exposure (Fig. S3). Synergistic activation of CRH^PVN^ and Hcrt^LH^ neurons during restraint stress is consistent with our observation of cFos staining as described in Fig. S1.

### Mild optogenetic stimulation of LH-projecting CRH^PVN^ neurons leads to insomnia

Since CRH^PVN^ neurons are sensitive to stressful stimuli, we hypothesized that the neuronal circuit connecting CRH^PVN^ and Hcrt^LH^ neurons drives hyperarousal in response to acute stress exposure. We thus conducted a series of optogenetic experiments in Hcrt::Cre (Fig. 3A), CRH::Cre (Fig. 3E), CRH::Cre-ATA3 mice (*33*) which have their Hcrt neurons ablated (Fig. 3I), and CRH::Cre-Cas9 mice with the *crh* gene disrupted using CRISPR-Cas9 technology (*34*) in CRH^PVN^ neurons (Fig. 3M). We used a mild optogenetic stimulation paradigm (i.e., 15 ms 473 nm blue light pulse at 0.1 Hz for 6 hours starting from the beginning of light phase) with neglectable effects on Hcrt^LH^ neurons in changing the amount of sleep/wakefulness (Fig. 3A-D&Q, Fig. S4A&E). Strikingly, this same optogenetic stimulation of CRH^PVN^ neurons labeled with ChR2-eYFP by AAV-Retro vectors injected to LH significantly increased the amount of wakefulness (^†^*P* < 0.0005, Fig. 3E-H&Q, Fig. S4B&F), here defined as insomnia/hyperarousal. This observation is consistent with earlier observation that i.c.v. injection of CRH causes long-lasting wakefulness (*35*). Interestingly, optogenetic stimulation of LH-projecting CRH^PVN^ neurons of the CRH::Cre-ATA3 mice without Hcrt neurons can also trigger a robust increase of wakefulness but the effect was significantly smaller when compared with the condition with intact Hcrt neurons (^†^*P* < 0.0005, Fig. 3I-L&Q, Fig. S4C&G). We further investigated whether CRH release was necessary for the optogenetically-induced arousal response. We generated viruses carrying sgRNAs to selectively disrupt the *crh* gene in CRH^PVN^ neurons of CRH::Cre-Cas9 mice (Fig. S5). Optogenetic stimulation of CRH^PVN^ neurons projecting to LH from *crh* gene-disrupted mice failed to increase wakefulness (Fig. 3M-P&Q, Fig. S4D&H) suggesting a critical role of CRH in activating downstream arousal promoting brain nuclei including the Hcrt^LH^ neurons (*36*).

**Fig. 3.**
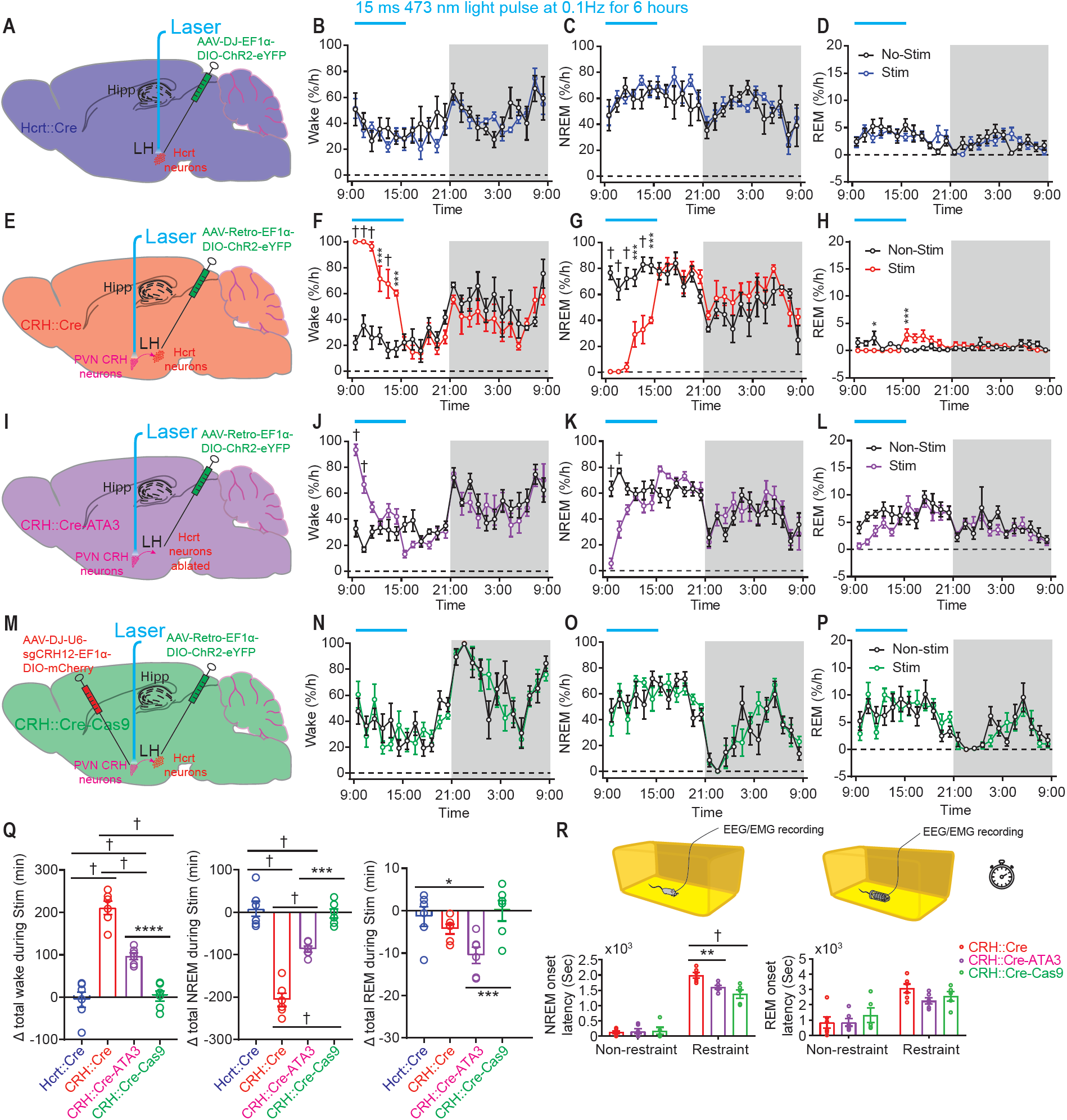
Optogenetic stimulation of CRH^**PVN**^ **neurons elicited strong arousal**. (**A**) Viral injection and fiber placement for optogenetic stimulation of Hcrt neurons in Hcrt::Cre mice. **(B-D)** Prolonged mild optogenetic stimulation of Hcrt^LH^ neurons had no significant effect on sleep/wake pattern. Hourly-based wake percentage (**B**, Group: F(1,10) = 4.539, *P* = 0.0590; Time: F(23, 230) = 3.808, ^†^*P* < 0.0001; Interaction F(23, 230) = 0.7973, *P* = 0.7334), hourly-based NREM sleep percentage (**C**, Group: F(1,10) = 4.791, *P* = 0.0534; Time: F(23, 230) = 3.556, ^†^*P* < 0.0001; Interaction F(23, 230) = 0.7164, *P* = 0.8271), hourly-based REM sleep percentage (**D**, Group: F(1,10) = 0.2327, *P* = 0.6399; Time: F(23, 230) = 3.293, ^†^*P*<0.0001; Interaction F(23, 230) = 1.268, *P* = 0.1909). (**E**) Viral injection and fiber placement for optogenetic stimulation of LH-projecting CRH^PVN^ neurons in CRH::Cre mice. **(F-H)** Prolonged mild optogenetic stimulation of CRH^PVN^ neurons elicited consolidated wakefulness. Hourly-based wake percentage (**F**, Group: F(1,10) = 28.68, ^†^*P* = 0.0003; Time: F(23, 230) = 7.163, ^†^*P* < 0.0001; Interaction F(23, 230) = 7.790, ^†^*P* < 0.0001), hourly-based NREM sleep percentage (**G**, Group: F(1,10) = 25.70, \*\*\*\**P* = 0.0005; Time: F(23, 230) = 7.143, ^†^*P* < 0.0001; Interaction F(23, 230) = 7.611, ^†^*P*<0.0001), hourly-based REM sleep percentage (**H**, Group: F(1,10) = 0.1275, *P* = 0.7285; Time: F(23, 230) = 1.340, *P* = 0.1431; Interaction F(23, 230) = 3.204, ^†^*P* < 0.0001). (**I**) Viral injection and fiber placement for optogenetic stimulation of CRH^PVN^ neurons projecting to LH in CRH::Cre-ATA3 mice. **(J-L)**, Prolonged mild optogenetic stimulation of CRH^PVN^ LH-projecting neurons significantly increased the amount of wakefulness in the absence of Hcrt neurons. Hourly-based wake percentage (**J**, Group: F(1,10) = 7.784, \**P* = 0.0191; Time: F(23, 230) = 9.536, ^†^*P* < 0.0001; Interaction F(23, 230) = 4.170, ^†^*P*<0.0001), hourly-based NREM sleep percentage (**K**, Group: F(1,10) = 1.517, *P* = 0.2463; Time: F(23, 230) = 8.722, ^†^*P* < 0.0001; Interaction F(23, 230) = 4.210, ^†^*P* < 0.0001), hourly-based REM sleep percentage (**L**, Group: F(1,10) = 2.899, *P* = 0.1194; Time: F(23, 230) = 5.629, ^†^*P* < 0.0001; Interaction F(23, 230) = 1.187, *P* = 0.2581). (**M**) Viral injection and fiber placement for optogenetic stimulation of LH-projecting CRH^PVN^ neurons with CRH gene interrupted. **(N-P)**, Disruption of CRH gene in CRH^PVN^ neurons compromised the strong wakefulness-promoting effect elicited by stimulating LH-projecting CRH^PVN^ neurons in CRH::Cre-Cas9 mice. Hourly-based wake percentage (**N**, Group: F(1,10) = 0.1892, *P* = 0.6644; Time: F(23, 230) = 14.30, ^†^*P* < 0.0001; Interaction F(23, 230) = 1.352, *P* = 0.1365), hourly-based NREM sleep percentage (**O**, Group: F(1,10) = 0.1215, *P* = 0.7346; Time: F(23, 230) = 14.42, ^†^*P* < 0.0001; Interaction F(23, 230) = 1.352, *P* = 0.1362), hourly-based REM sleep percentage (**P**, Group: F(1,10) = 0.003993, *P* = 0.9509; Time: F(23, 230) = 8.442, ^†^*P* < 0.0001; Interaction F(23, 230)=1.136, *P*=0.3073). (**Q**) Prolonged mild optogenetic stimulation of LH-projecting CRH^PVN^ neurons but not Hcrt neurons increases the wake amount. Absence of Hcrt neurons or CRISPR-Cas9-mediated disruption of CRH gene in CRH^PVN^ neurons compromised this effect (left, F(3, 20) = 51.44, ^†^*P* < 0.0001; middle, F(3, 20) = 54.20, ^†^*P* < 0.0001; right, F(3, 20) = 5.355, \*\**P* = 0.0072;). (**R**) Absence of Hcrt neurons or disruption of CRH gene in CRH^PVN^ neurons significantly reduced the latency to NREM sleep onset after a 10 min restraint stress session (NREM Non-restraint: F(2, 14) = 0.08136, *P* = 0.9223; NREM Restraint: F(2, 14) = 10.17, \*\*\**P* = 0.0019; REM Non-restraint: F(2, 14) = 0.5333, *P*=0.5891; REM Restraint: F(2, 14) = 2.529, *P* = 0.1154; **B-D, F-H, J-L, N-P**: linear mixed-effects model followed by Šidák’s multiple comparisons, dark phase indicated by gray shielding; **Q**&**R**: one-way ANOVA followed by Šidák’s multiple comparisons; \**P* < 0.05, \*\**P* < 0.01, \*\*\**P* < 0.005, \*\*\*\**P* < 0.001, ^†^*P* < 0.0005; n = 6 mice for each group except for n = 5 for CRH::Cre-Cas9 in panel **r**).

### *Crh* gene disruption compromises hyperarousal/insomnia caused by restraint stress

Given the observation of robust wakefulness upon optogenetic activation of LH-Projecting CRH^PVN^ neurons (Fig. 3 & Fig. S4), we then evaluated whether the CRH^PVN^-Hcrt^LH^ pathway is relevant to hyperarousal/insomnia under stress. We submitted three genotypes of mice to restraint stress with intact or disrupted CRH^PVN^-Hcrt^LH^ neuronal circuitry. We monitored the EEG/EMG following a 10 min restraint session (*22*) in CRH::Cre mice, CRH::Cre-ATA3 mice with a genetic ablation of Hcrt neurons, and CRH::Cre-Cas9 mice with *crh* gene disrupted (Fig. S5) bilaterally in PVN (Fig. 3R). A 10 min restraint stress session significantly delayed the NREM and REM sleep onset. Interestingly, either genetic ablation of Hcrt neurons or CRISPR-Cas9-mediated disruption of the *crh* gene in CRH^PVN^ neurons significantly reduced the latency to NREM sleep onset compared to the mice with intact Hcrt neurons and *crh* gene, suggesting that these neurons play a synergistic role in stress-induced insomnia/hyperarousal.

### Optogenetic activation of CRH^PVN^ neurons causes immunosuppression

As activation of stress circuitry drives changes in whole-body physiology (i.e., ‘fight or flight’ response), we investigated the immunomodulatory effects of CRH neuron stimulation using single-cell mass cytometry by time of flight (CyTOF) (*37*). We assessed changes in the distribution of circulating immune cells (Fig. 4) as well as alterations in intracellular signaling cascades (Fig. S6) in response to a single session of CRH^PVN^ optogenetic stimulation. CRH neuron stimulation resulted in dramatic alterations in peripheral immune cell distribution (Fig. 4B). Specifically, we observed a reduction in the frequencies of NK cells, MHCII+ dendritic cells, inflammatory (Ly6C^Hi^), patrolling (Ly6C^low^) monocytes, B cells (total), Th1 TBet+ T cells and CD4+ T cells (total) in response to optogenetic stimulation (Fig. 4C&D). These findings indicate that CRH neuronal stimulation altered components of both the innate and adaptive immune system. CRH neuron stimulation may alter immune cell distribution through several mechanisms including alteration of intracellular signaling responses implicated in the mobilization, adhesion, proliferation and survival of these cells (*38*). We next investigated the effect of CRH neuron stimulation on the activity of key signaling pathways implicated in the immune response to stress, including elements of the NFκB, MAPK and JAK/STAT signaling pathways. CRH neuron stimulation promoted expression of IκB in nearly all cell types examined (Fig. 4B left panel & Fig. S6), consistent with the known inhibitory effects of glucocorticoids on NFκB signaling (*39, 40*). Additionally, optogenetic stimulation down regulated pSTAT6 protein expression in many immune cell subsets, including B cells, CD4+ and CD8+ T cells, and NK cells, consistent with prior observations of direct interactions between the glucocorticoid receptor and STAT6 (*41*). The inhibition of NFκB and STAT6 signaling responses in T cells and monocyte subsets, as well as the observed reductions in the numbers of circulating CD4+ T cells, B220+ B cells, NK cells, and monocyte subsets suggests that CRH neuronal stimulation promoted widespread immunosuppression consistent with known glucocorticoid responses (*42*). Our observation of central optogenetically-induced immunosuppression is consistent with an enhancement of glucocorticoid signaling by physical restraint stress as reported earlier (*43*). Therefore, we identified a neural substrate for stress-induced immunosuppression.

**Fig. 4.**
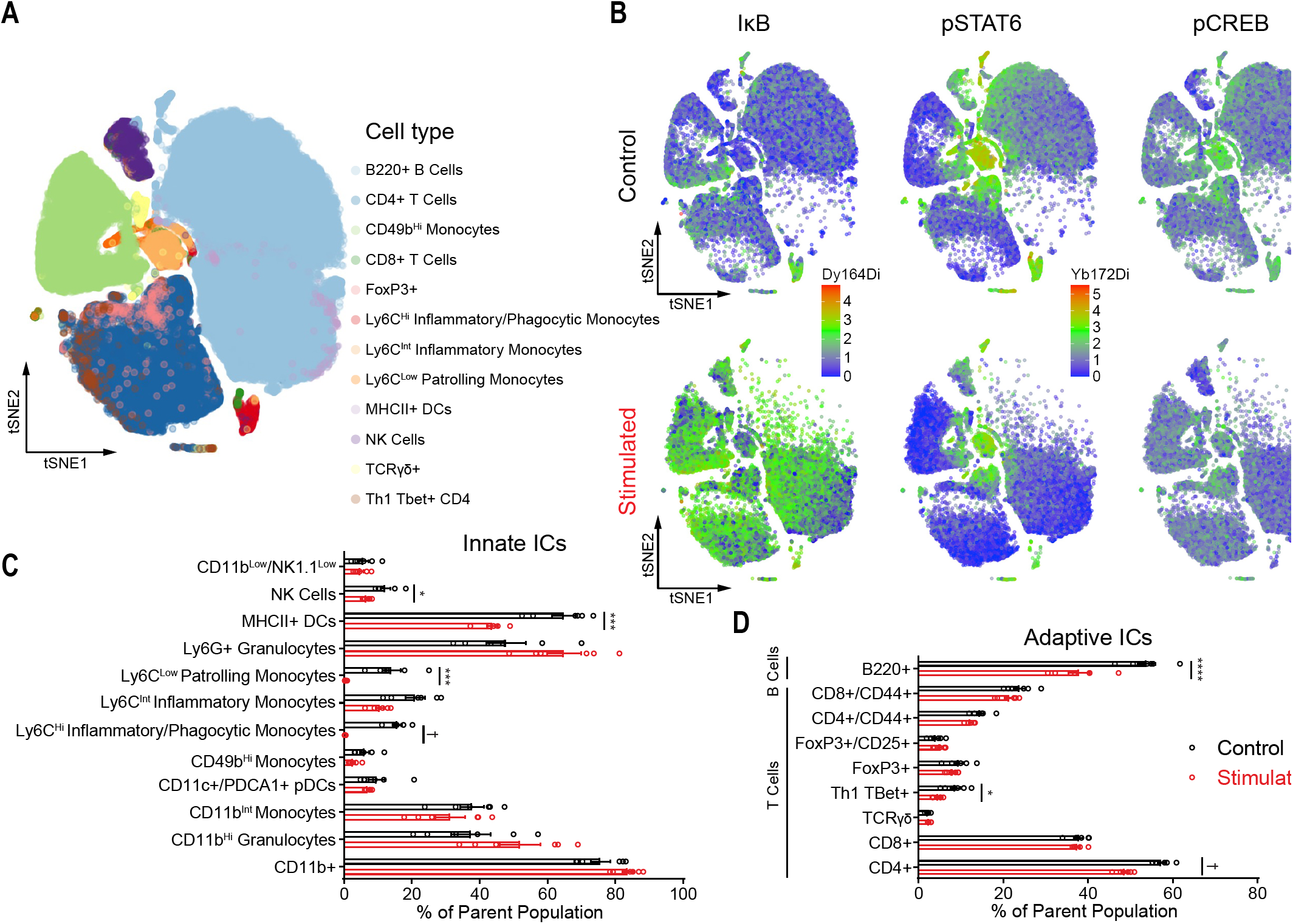
CRH^**PVN**^ **activation drives changes in systemic immunity consistent with immunosuppression**. (**A**) Cell subpopulation distribution in the pooled dataset based on all samples. (**B**) Composite tSNE plots of intracellular protein expression from control and optogenetically stimulated mice for IκB (left), pSTAT6 (middle), and pCREB (right). (**C**) Distribution of circulating innate immune cells (ICs) in response to CRH neuronal stimulation including NK1.1+ NK cells (control mean: 12.17%; stimulation mean: 6.665%, t ratio = 3.617, adjusted \**P* = 0.041632), MHCII+ dendritic cells (control mean: 64.83%; stimulation mean: 43.74%, t ratio = 5.496, adjusted \*\*\**P* = 0.002891), Ly6C^Low^ patrolling monocytes (control mean: 14.01%; stimulation mean: 0.4224, t ratio = 2.688, adjusted \*\*\**P* = 0.004937), and Ly6C^Hi^ Inflammatory/Phagocytic monocytes (control mean: 15.72%; stimulation mean: 0.2877%; t ratio = 12.62, adjusted ^†^*P* = 0.000002). (**D**) Distribution of adaptive ICs in circulation following CRH neuronal stimulation. B lymphocytes (B220+) were decreased (control mean: 53.83%; stimulation mean: 37.94%; t ratio = 4.875, adjusted \*\*\*\**P* = 0.000647). T lymphocyte subsets altered include Th1 TBet+ cells (control mean: 8.733%; stimulation mean: 4.693%; t ratio = 3.365, adjusted \**P* = 0.049209) and CD4+ T cells (control mean: 57.35%; stimulation mean: 48.7%; t ratio = 7.232, ^†^*P* = 0.000225) (n = 6 mice/group, error bars represent SEM, Holm-Šidák test, \**P* < 0.05, \*\**P* < 0.01, \*\*\**P* < 0.005, \*\*\*\**P* < 0.001, ^†^*P* < 0.0005).

## DISCUSSION

In this study, we find that the CRH^PVN^-Hcrt^LH^ pathway plays a crucial role underlying stress-induced insomnia/hyperarousal and systemic immunosuppression. We show that CRH^PVN^ neurons are sufficient but not necessary to trigger Hcrt^LH^ neuronal activity. We further demonstrate that optogenetic activation of CRH^PVN^ neurons projecting to the LH elicits long-lasting wakefulness mimicking insomnia/hyperarousal induced by stress exposure. Importantly, we find that CRH release from PVN neurons is necessary for optogenetically-evoked hyperarousal and restraint stress-induced insomnia. These observations are in line with earlier work showing stress induces CRH release in PVN (*44*), and knockdown of CRH attenuates stress-induced social avoidance (*45*). In addition, ablation of Hcrt neurons partially reduced the amount of wakefulness elicited by optogenetically stimulating CRH^PVN^ neurons projecting to LH and decreased the latency to NREM sleep onset following restraint stress. Our work highlights the CRH^PVN^ signaling pathway is a prominent target for the treatment of insomnia/hyperarousal and stress-induced changes in systemic physiology.

### Optical dissection of CRH^PVN^-Hcrt^LH^ circuitry in insomnia induced by stress

cFos staining of brain slices following restrain stress informed us the hyperactivity of CRH^PVN^ and Hcrt^LH^ neurons are potentially implicated in insomnia pathology. Many brain nuclei including hypocretinergic (*6*), aminergic (*8-10*), and basal forebrain cholinergic neurons (*12, 46*) are capable of switching the animals from sleep-to-wake. However, the optogenetic stimulation paradigm used in these experiments are usually consist of multiple light pulses at frequencies mimicking or above their natural neuronal firing rates, and insomnia/hyperarousal has not been reported following optogenetic stimulation of these brain nuclei. In this study, we used a mild stimulation paradigm (10 mW, 15 ms pulse at 0.1 Hz for 6 hours) which is insufficient to increase wakefulness amount in Hcrt::Cre mice. Surprisingly, this mild stimulation of CRH^PVN^ neurons projecting to LH where Hcrt neurons are located dramatically elevated the arousal level mimicking insomnia under stress confirming earlier observation showing single central administration of CRH causes long-lasting wakefulness (*35*). We further investigated the specific role of Hcrt neurons and *crh* gene in PVN in the same stimulation scenario. Interestingly, genetic ablation of Hcrt neurons significantly reduced the wakefulness amount upon optogenetic stimulation of LH projecting CRH^PVN^ neurons and CRISPR-Cas9-mediated interruption of *crh* gene completely abolished the hyperarousal. These observations highlight the CRH hormone plays an essential role in stress-induced insomnia/hyperarousal. The partial compromise of hyperarousal by genetic ablation of Hcrt neurons indicates CRH exerts arousal-promoting effect through other downstream neurons besides Hcrt^LH^ neurons. Fiber photometry recording revealed CRH^PVN^ neurons are silent during natural sleep-to-wake transitions, but highly active under stress. The concomitant Hcrt^LH^ neuronal activity under stress suggests CRH^PVN^ neurons recruit Hcrt neurons to mediate hyperarousal. Response of Hcrt^LH^ neurons to neutral/positive stimuli indicates Hcrt neurons could be engaged in various conditions requiring vigilance independent of valence.

### Neural substrates of stress-induced peripheral immunosuppression

Stress-induced insomnia is often associated with deficits in immune function in humans (*47*). However, the neural substrates linking stress, arousal, and immunity are poorly understood. As we discussed above, CRH neurons in the paraventricular nucleus of the hypothalamus serve as a key node in the behavioral and physiological stress response in part via the promotion of downstream glucocorticoid secretion from the adrenal cortex. We identified the CRH^PVN^-Hcrt^LH^ circuit as a critical regulator coupling stress to arousal and protracted insomnia. Our observations that stimulation of CRH^PVN^ soma resulted in dynamic changes in immune cell distribution and functional responses (e.g., reduction in circulating CD4+ T cells and increases in IκB expression) is consistent with known immunosuppressive effects of glucocorticoids. However, as acute and chronic stress have frequently opposite effects on circulating immune cell function, we were not expecting to observe such a strong immunosuppressive phenotype in response to just one session of optogenetic stimulation (lasting 1 hour). Further studies are needed to understand how different stimulation paradigms influence systemic and tissue-specific immune cell populations.

An alternative explanation (beyond glucocorticoid upregulation) for the changes we observed may depend on CRH^PVN^-mediated activation of the sympathetic nervous system. Indeed, CRH neurons in the paraventricular nuclei drives downstream sympathetic activation via projections to the brainstem nucleus of the solitary tract (*48*). Post-ganglionic sympathetic nerves have well known immunomodulatory effects, driven largely by the effect of their neurotransmitter product, norepinephrine (NE) (*48*). In early studies, the effects of CRH on the immune system appeared to be dependent on the sympathetic nervous system, as chemical sympathectomy and pharmacological blockade of beta-adrenergic receptors reversed the observed effect of CRH on immune function (*49*). If the immunosuppressive phenotype we observed depends on sympathetic activation, then beta adrenergic blockade may inadvertently prevent immune dysfunction associated with stress and insomnia (e.g., as in (*50*)).

Together, we reveal that the CRH^PVN^-Hcrt^LH^ pathway plays an essential role in hyperarousal via optical interrogation of sleep and CyTOF analysis of systemic immunity, therefore identifying a shared neural substrate for insomnia and immunosuppression under stress.

## MATERIALS AND METHODS

### Animals

Experiments were performed following the protocols approved by the Stanford University Animal Care and Use Committee in accordance with the *National Institutes of Health Guide for the Care and Use of Laboratory Animals*. Discomfort, distress, or pain were minimized with anesthesia and analgesic medications. Young adult (3-6 months) male mice were used in this study. Mice were housed in a temperature- and humidity- controlled animal facility with a 12-h/12-h light/dark cycle (9:00 am light on, 9:00 pm light off). Standard laboratory mouse food pellet and water were available *ad libitum*. Hcrt-IRES-Cre knock-in heterozygotes (Hcrt::Cre)(*14*), orexin/ataxin-3 mice (ATA3)(*33*), and CRH::Cre (B6(Cg)-Crh^tm1(cre)Zjh^/J, JAX Stock No: 012704), AVP::Cre (B6.Cg-Avp^tm1.1(cre)Hze^/J, JAX Stock No: 023530), OXT::Cre (B6;129S- Oxt^tm1.1(cre)Dolsn^/J, JAX Stock No: 024234) were backcrossed onto C57BL/J backgrounds. CRH::Cre-ATA3 mice were generated by crossing CRH::Cre heterozygotes with ATA3 heterozygotes. CRH::Cre-Cas9 mice were generated by crossing CRH::Cre heterozygotes with Cas9 homozygotes (B6J.129(B6N)- Gt(ROSA)26Sor^tm1(CAG-cas9*,-EGFP)Fezh^/J, JAX Stock No:026175).

### EEG/EMG electrode fabrication and implantation

Mini-screw (US Micro Screw) was soldered to one tip of an insulated mini wire with two tips exposed, and the other tip of the mini-wire was soldered to a golden pin aligned in an electrode socket. A mini-ring was made on one side of an insulated mini-wire with the other side soldered to a separate golden pin in the electrode socket. Each electrode socket contains 4 channels with 2 mini-screw channels for EEG recording and 2 mini-ring channels for EMG recording. The resistance of all the channels were controlled with a digital Multimeter (Fluke) to be lower than 1.5 Ω for ideal conductance. Mice were mounted onto an animal stereotaxic frame (David Kopf Instruments) under anesthesia with a mixture of Ketamine (100 mg/kg) and Xylzine (20 mg/kg). Two mini-screws were placed in the skull above the frontal (AP: -2 mm; ML: ± 1 mm) and temporal (AP: 3 mm, ML: ± 2.5 mm) cortices for sampling EEG signals and two mini-rings were placed in the neck muscles for EMG signal acquisition. Electrode sockets were secured with meta-bond and dental acrylic on skull for recording in freely moving mice. After surgery, mice were administered with buprenorphine-SR (0.5 mg/kg 30 min before and 24-48 hours post surgery, SC) for pain relief.

### EEG/EMG recording and analysis

EEG/EMG signals were sampled at 256 Hz with VitalRecorder (Kissei Comtec Co.) following amplification through an amplifier multiple channels (Grass Instruments). The bandpass was set between 0.1-120 Hz. Raw EEG/EMG data were exported to Matlab (MathWorks, Natick, MA, USA) and analyzed with custom scripts and Matlab built-in tools based on described criteria (*6*) to determine behavioral states. For optogenetic and fiber photometry recording experiments, simultaneous EEG/EMG signals were recorded to determine behavioral states. The raw EEG/EMG signals with/without GCaMP6f traces were imported to Matlab for offline analysis.

### Virus injection and fiber optic implantation

#### Monosynaptic tracing of inputs to Hcrt^*LH*^ *neurons*

To determine the cell type in the PVN with direct innervation of Hcrt^LH^ neurons, we used the EnvA-pseudotyped glycoprotein (G)-deleted rabies virus (EnvA + RV*d*G) tracing strategy (*26*). Firstly, a mixture of AAV-5 EF1α-Flex TVA-mCherry (AAV67, lot # 388b, GC: 2.5 × 10^12^, Stanford virus core) and AAV- 8(Y733F)-CAG Flex RabiesG (AAV-59, Lot#1489, GC 5.0 × 10^13^, Stanford virus core) was injected to LH of young (3-6 month) male Hcrt::Cre mice to target the Hcrt field (AP: −1.35 mm, ML: ±0.95 mm, DV: −5.15 mm) under anesthesia and analgesic as described above. Two weeks later, the same mice were injected with EnvA G-deleted Rabies GFP (GC: 6.04 × 10^7^, Salk Institute Gene Transfer Targeting and Therapeutics Core) at the coordinates same with the first injection. One week after the second injection, colchicine (1.0 μl of 7 μg/ml solution per hemisphere, for PVN CRH/AVP/OXT neuropeptide immunostaining) was injected to the lateral ventricles (AP: –0.65 mm, ML: ± 0.95 mm, DV: –2.55 mm). Then mice were euthanized and perfused 24-48h later for immunostaining (Fig. 1).

#### AAV-Retro tracing of LH upstream neurons

PVN is known as a hub containing intermingled CRH, AVP and OXT neurons (*23*). To further validate the LH receives neuronal projections from PVN, we injected AAV-Retro-EF1α-DIO-hChR2(C128S/D156A)- eYFP (AAV-Retro, Stanford Virus Core, 3.3 × 10^13^ gc/ml Lot# 3908) to the LH containing Hcrt field (AP: −1.35 mm, ML: ±0.95 mm, DV: −5.15 mm) of young male CRH::Cre, AVP::Cre, OXT::Cre mice respectively under anesthesia and analgesic as described above. Three weeks after injection, colchicine (1.0 μl of 7 μg/ml solution per hemisphere, for PVN CRH/AVP/OXT neuropeptide immunostaining) was injected to the lateral ventricles (AP: –0.65 mm, ML: ± 0.95 mm, DV: –2.55 mm). Then mice were euthanized and perfused 24-48h later for immunostaining (Fig. S2).

#### Fiber photometry

For fiber photometry recording from Hcrt^LH^ (also known as orexin(*51, 52*)) neurons in Hcrt::Cre mice, 0.3 μl AAV vectors carrying genes encoding GCaMP6f (AAV-DJ-EF1α-DIO-GCaMP6f, 1.1 × 10^12^ gc/ml, Stanford Virus Core, lot# 3725) were delivered to L/R LH (AP: −1.35 mm, ML: ±0.95 mm, DV: −5.15 mm) of young male Hcrt::Cre mice with a 5 μl Hamilton micro-syringe, and a glass fiber (400 μm core diameter, Doric Lenses) was implanted with the tip at the injection site for GCaMP6f signal acquisition later on. For recording from CRH^PVN^ neurons, same virus was injected to L/R PVN (AP: −0.90 mm, ML: ±0.30 mm, DV: −4.66 mm) and glass fiber was placed at the same site of young CRH::Cre mice. EEG and EMG electrodes were implanted after securing the fiber optic with meta-bond and dental acrylic. Mice were housed in their home cages to recover for at least 2 weeks to get sufficient viral expression before connection to the EEG/EMG recording cables and fiber photometry recording patch cord.

#### Optogenetic experiments

For optogenetic experiments, 0.3 μl AAV-DJ-EF1α-DIO-hChR2 (H134R)-eYFP viruses (ChR2-eYFP, 6.5 × 10^12^, Stanford Virus Core, Lot# 4176) was delivered to LH (AP: −1.35 mm, ML: ±0.95 mm, DV: −5.15 mm) of anesthetized young (3-6 months) male Hcrt::Cre mice according to stereotaxic coordinates determined on Kopf stereotaxic frame with a 5 μl Hamilton microsyringe. A glass fiber (200 μm core diameter, Doric Lenses, Franquet, Québec, Canada) was implanted with the tip right above the injection site for optogenetic stimulations later on. For stimulating CRH^PVN^ neurons projecting to LH, AAV-Retro-EF1α-DIO- hChR2(C128S/D156A)-eYFP (AAV-Retro, Stanford Virus Core, 3.3 × 10^13^ gc/ml Lot# 3908) was injected to LH with the same coordinates for Hcrt::Cre mice. A glass fiber (200 μm core diameter, Doric Lenses, Franquet, Québec, Canada) was implanted with the tip right above the CRH^PVN^ neurons (AP: −0.90 mm, ML: ±0.30 mm, DV: −4.3 mm) for optogenetic stimulations of LH-projecting CRH^PVN^ neurons later on. After fixation of glass fiber and EEG/EMG electrode implantation, mice were allowed to recover for at least 2 weeks to get sufficient viral expression before connecting to the EEG/EMG recording cables and optical stimulation patch cord.

### CRISPR-Cas9-medaited *crh* gene disruption in CRH::Cre-Cas9 mice

*Crh* gene target sites for CRISPR/Cas9 were designed by CHOPCHOP (http://chopchop.cbu.uib.no)(*53*). The target sequences were as follows; sgCRH1: 5′-ATGCGGATCAGAACCGGCTG-3′, sgCRH2: 5′- CAACTCCACGCCCCTCACCG-3′. Oligonucleotides encoding guide sequences are purchased from Integrated DNA Technologies (IDT) and cloned individually into BbsI fragment of pX458 (Addgene plasmid 48138(*54*)). U6-sgCRH1and U6-sgCRH2 fragments were PCR-amplified, respectively using pX458-sgCRH as a template. Amplified fragments were cloned tandemly into MluI-digested pAAV EF1α DIO mCherry (Addgene plasmid 20299) by Gibson assembly method. The primers used were as follows; Gibson1-F: 5′- TAGGGGTTCCTGCGGCCGCAGAGGGCCTATTTCCCATG-3′, Gibson1-R: 5′- ATAGGCCCTCGGTACCAAAAATCTCGCC-3′, Gibson2-F: 5′- TTTTTCTAGAGAGGGCCTATTTCCCATG-3′, Gibson2-R: 5′- ATCCATCTTTGCAAAGCTTAAAAAATCTCGCCAACAAGTTG-3′. AAV construct carrying non-targeting guide sequences were used as control (*34*). pAAV U6 sgCRH1-U6 sgCRH2 EF1α DIO mCherry, and pAAV U6 sgControl-U6 sgControl EF1α DIO mCherry were packaged into AAV-DJ by the gene vector and virus core at Stanford University. For *crh* gene interruption validation, 0.6 μl AAV-DJ-U6- sgControl- EF1α-DIO-mCherry (5 × 10^12^ gc/ml Lot# 4354) / AAV-DJ-U6-sgCRH12-EF1α-DIO-mCherry viruses (3 × 10^12^ gc/ml Lot#4553) were injected to PVN (AP: −0.90 mm, ML: ±0.30 mm, DV: −4.5 + -4.8 mm, 0.3 μl each depth) unilaterally of the CRH::Cre-Cas9 for *crh* gene disruption validation (Fig. S5). Three weeks after the virus injection, colchicine (1.0 μl of 7 μg/ml solution per hemisphere, for CRH neuropeptide immunostaining) was injected to the lateral ventricles (AP: –0.65 mm, ML: ± 0.95 mm, DV: – 2.55 mm). Then mice were euthanized and perfused 24-48h later for immunostaining. For optogenetic stimulation of LH-projecting CRH^PVN^ neurons with *crh* gene interrupted, 0.6 μl AAV-DJ-U6-sgCRH12- EF1α-DIO-mCherry viruses were injected to the L/R PVN (−0.90 mm, ML: ±0.30 mm, DV: −4.5 + -4.8 mm, 0.3 μl each depth) unilaterally of CRH::Cre-Cas9 mice under anesthesia, and 0.3 μl AAV-Retro- EF1α-DIO-hChR2(C128S/D156A)-eYFP viruses were injected to the LH (AP: −1.35 mm, ML: ±0.95 mm, DV: −5.15 mm) of the same hemisphere. Afterwards, a glass fiber (200 μm core diameter, Doric Lenses) targeting PVN infected by viral vectors and EEG/EMG electrodes were implanted for optogenetic stimulation of *crh* gene ablated CRH^PVN^ neurons projecting to LH later on (Fig. 3M). For the restraint stress on sleep experiment (Fig. 3R), 0.6 μl AAV-DJ-U6-sgCRH12-EF1α-DIO-mCherry viruses were injected to the PVN (−0.90 mm, ML: ±0.30 mm, DV: −4.5 + -4.8 mm, 0.3 μl each depth) bilaterally of CRH::Cre-Cas9 mice under anesthesia, and EEG/EMG electrodes were implanted for sleep pattern recording.

### Restraint stress and sleep recording

A well-established non-invasive restraint paradigm was used to generate acute stress in mice (*22*). For the cFos immunostaining experiment (Fig. S1), 9 young adult male WT mice (3-5 months) were individually placed head-first into a well-ventilated 50-ml Falcon conical tube with a narrow open window on top (Fig. 3R top panel) placed in their home cages. After a 10 min restraint session, mice were immediately released from the restraint tubes. Mice were randomly separated into 3 groups, and were perfused at 20 min (n = 3), 80 min (n = 3), and 140 min (n = 3) after the restraint stress termination. For evaluating the impact of restraint stress on latency to sleep onset, young adult male mice (3-6 months, CRH::Cre, CRH::Cre-ATA3, CRH::Cre-Cas9 infused with *crh* gene disrupted bilaterally in CRH^PVN^ neurons) with EEG/EMG implants (for sleep pattern monitoring) were exposed to the restraint tubes with a narrow window on top for EEG/EMG cable sliding (Fig. 3R top panel). During the entire restraint procedure, mice were in a natural body position without physical harm in their home cages. The restraint stress lasted for 10 min at the beginning of light phase when mice have a strong drive for sleep. EEG/EMG signals were recorded starting from the onset of restraint stress. Mice were released from the restraint tubes immediately after a 10 min restraint session. Mice were assigned to non-restraint and restraint paradigms according to a counterbalanced crossover design.

### Salient stimuli

GCaMP6f signal was recorded from CRH^PVN^ neurons in CRH::Cre mice and Hcrt^LH^ neurons in Hcrt::Cre mice housed individually with 1 week habituation to fiber patch cord. GCaMP6f signal was sampled for 4 minutes each session with 2 minutes as baseline and 2 minutes exposure to salient stimuli (Fig. S3) including a 15 ml novel Falcon tube (novelty), novel male mouse, novel female mouse, high fat diet (in a small petri dish), rat bedding (in a small petri dish), restraint (mouse placed in a well-ventilated 50ml Falcon conical tube with a narrow window for fiber patch cord sliding), TMT (a small piece of filtration paper with 5 μl TMT in a small petri dish), grabbing. For the elevated plus (EPM) maze open arm exposure, mouse was transferred from the home cage onto the open arm of the EPM with the entrance to EPM center blocked. Salient stimuli were removed immediately from the mice after 2 minutes. Introduction of salient stimuli to each individual mouse was randomly shuffled.

### Fiber photometry signal acquisition and analysis

Following recovery and habituation to EEG/EMG cable lead and fiber optic patch cord (400 μm core diameter, Doric Lenses), mice expressing Cre-dependent GCaMP6f were connected to EEG/EMG recording setup and custom-built fiber photometry setup (*55*) respectively. Briefly, a 470 nm LED (M470D3, Thorlabs, NJ, USA) was sinusoidally modulated at 211 Hz and passed through a GFP excitation filter followed by a dichroic mirror (MD 498, ThorLabs) for reflection. The light stream was sent through a high NA (0.48), large core (400 μm) optical fiber patch cord (Doric Lenses, Québec, Canada), which was connected with a zirconia connector (Doric Lenses, Québec, Canada) to the dental acrylic-secured fiber optic implant (0.48NA, 400 μm, Doric Lenses, Québec, Canada) with tip above the injection site targeting Hcrt neurons. Separately, a 405 nm LED was modulated at 531 Hz and filtered by a 405 nm bandpass filter and sent through the optical fiber patch cord to mouse brain to evoke reference fluorescence, which is independent of Ca^2+^ release. GCaMP6f fluorescence and reference fluorescence were sampled by the same fiber patch cord through a GFP emission filter (MF525-39, ThorLabs), and center-aligned to a photodetector (Model 2151, Newport, Irvine, CA, USA) with a lens (LA1540-A, ThorLabs). The analog signals were amplified by two lock-in amplifiers for GFP fluorescence and reference fluorescence respectively (30 ms time constant, model SR380, Stanford Research Systems, Sunnyvale, CA, USA). Matlab-based custom software was used to control the LEDs and sample both the GFP fluorescence and reference fluorescence through a multifunction data acquisition device (National Instruments, Austin, TX, USA) with 256 Hz sampling frequency in a real-time manner. ΔF/F was obtained by subtracting the reference fluorescence signal from the 470 nm excited GFP emission signal to remove the system interference. GCaMP6f from CRH^PVN^ or Hcrt^LH^ neurons and simultaneous EEG/EMG recordings were conducted between 14:00-18:00 during the light phase. CaMP6f signals were staged to different behavioral state transitions based on the simultaneously recorded EEG/EMG signals (Fig. 2). The behavioral state transition category include NREM to brief wake, NREM to REM, NREM to long wake, REM to brief wake, REM to long wake and Wake to NREM. Traces for each behavioral state transitions were plotted with transition time point defined as 0 for Hcrt::Cre and CRH::Cre mice. For GCaMP recording during salient stimuli challenges, time of introduction of the stimuli to the mice was defined as time 0. A Z score was calculated by subtracting the mean value of GCaMP6f after time 0 from the mean value of GCaMP6f trace prior to time 0.

### Optogenetic stimulation

After recovery, mice injected with viruses expressing Cre-dependent ChR2-eYFP were connected to EEG/EMG recording cables and fiber optic patch cord (200 μm core diameter, Doric Lenses) for one week acclimation in cages with open top which allows mice to move freely. Following acclimation, EEG and EMG were recorded for 24 hours covering an entire light/dark cycle with 15 ms 10 mW (LaserGlow, calibrated with Thorlabs light meter) 473 nm light pulse optogenetic stimulation at 0.1 Hz for 6 hours starting from the beginning of light phase. Control and stimulation were scheduled according to a counterbalanced crossover design.

### Histology

For *in vivo* experiments, upon accomplishment of recordings, mice were perfused under anesthesia (100 mg/kg Ketamine and 20 mg/kg Xylazine mixture) with ice-cold PBS to verify the viral expression and glass fiber placement. Brains were rapidly extracted, postfixed with 4% PFA at 4 °C overnight, and equilibrated in 30% sucrose in PBS containing 0.1% NaN3. Then, brains were sectioned at -22 °C with a cryostat (Leica Microsystems) at a thickness of 35 µm. Slices were collected from anterior to posterior consecutively to 24- well plates containing PBS with 0.1% NaN3, covered with aluminum foil, and stored at 4 °C until immunostaining and imaging. Primary antibody against OX-A/Hcrt-1 (SC-8070, Lot#A2915, Goat polyclonal IgG) was purchased from Santa Cruz Biotechnology, INC. Primary polyclonal antibody against cFos (Host: Rabbit, Catalog # 26209, Lot # 1714001), primary polyclonal antibody against CRH (Host: Rabbit, Catalog # 20084, Lot # 533038), primary polyclonal antibody against AVP (Host: Rabbit, Catalog # 20069, Lot # 1911001), and primary polyclonal antibody against OXT (Host: Rabbit, Catalog # 20068, Lot # 1607001), were purchased from ImmunoStar. Secondary antibodies Alexa Fluor 594 Donkey anti-Goat IgG (H+L, Catalog # A-11058), Alexa Fluor 594 Donkey anti-Rabbit IgG (H+L, Catalog # A32754), Alexa Fluor 488 Donkey anti-Rabbit IgG (H+L, Catalog # A-21206), Alexa Fluor 647 Donkey anti-Rabbit IgG (H+L, Catalog # A-31573) were purchased from Invitrogen (Manufacturer: Life Technologies). For the WT mice used for restraint stress, sections around PVN and LH were washed in PBS for 5 minutes, 3 times and incubated in a blocking solution of PBS with 0.3% Triton X-100 (PBST) and 4% bovine serum albumin (BSA) for 1 hour. Following that, CRH primary antibody and OX-A/Hcrt-1 primary antibody were added to the wells containing slices around PVN and wells containing LH slices respectively, and cFos primary antibody was added to both PVN slice containing wells and LH slice containing wells with blocking solution (1:800) overnight. On the second day, sections were washed in PBS for 5 minutes, 3 times, and incubated in blocking buffer for 2 hours. After blocking, secondary antibody was added to the blocking buffer for 2 hours (dilution 1:800). After 3 times of 5-min PBS washing, brain sections were mounted onto gelatin-coated slides, covered with Fluoroshield containing DAPI Mounting media (Sigma-Aldrich, F6057) and cover glass for imaging with wild field microscope (Zeiss AxioImager, Germany) for entire section or LSM710 Confocal Microscope for enlarged visualization (Zeiss, Germany).

### CyTOF

CRH::Cre mice were equipped with EEG/EMG electrodes as described previously. AAVs encoding the optogenetic actuator ChR2 (AAV-DJ-DIO-EF1α-ChR2-eYFP; Lot # 5227; 4.12 × 10^12^ gc/ml) were infused into the paraventricular nucleus (unilateral; 0.4 μl; -0.90 mm AP, ± 0.25 mm ML, -4.85 mm DV (injection), -4.6 mm DV (optic fiber)) over the course of 3 min followed by 10 min for viral distribution to the tissue. Then, a fiber optic implant (200 µm diameter, 5.5 mm length, Doric Lenses, Inc.) was slowly lowered and cemented in place above the PVN. Mice were allowed to recover for at least 3 weeks prior to starting the experiment.

Following recovery, 15 ms 10 mW (LaserGlow, calibrated with Thorlabs light meter) 473 nm light pulses at 10 Hz (10 sec ON, 20 sec OFF) were delivered to the PVN for 1 hour, and then whole blood was collected 3 hours following the start of stimulation using microcapillary tubes (retro-orbital bleed; approximately 350 μl blood). Blood was immediately stabilized using a protein stabilization reagent, allowed to sit for 10 minutes, and then placed at -80 °C in cryovials until further processing. Red blood cells (RBCs) were lysed using 1x Thaw-lyse buffer (Smart Tube Inc., San Carlos, CA) in ddH_2_ O. After lysing, cells were pelleted at 600× g for 5 min at room temperature. RBC depleted samples were barcoded using palladium isotopes. In brief, cells were washed with cell staining media (CSM), PBS, and PBS containing 0.02% saponin and then placed into a deep well block. Unique barcoding reagents were added to each well of the block containing cells from each mouse and incubated for 15 minutes at room temperature. After incubation, cells were washed with CSM and then resuspended via brief vortexing. Then, all samples were pooled into a single fluorescence-activated cell sorting (FACS) tube for the next staining step. After washing and pelleting the pooled cells, 150 μl of Fc block was added and cells were incubated at room temperature (RT) with gentle shaking (600 rpm) for 10 minutes. Then, 300 μl of the surface antibody mix (see **Table. S1**) was added and cells were incubated at RT for 30 minutes with gentle shaking. Then, cells were washed, pelleted, and the supernatant was collected and discarded. To permeabilize cells for intra-cellular staining, 700 μl of MeOH was added to the tube and cells were incubated at 4 °C for 10 minutes. Then, cells were washed with PBS and CSM, followed by incubation with the intracellular antibody mix (see **Table. S1**) for 30 minutes at RT with gentle shaking (600 rpm). Cells were washed and then a DNA intercalator (DVS # Inter-1X-natir) was added in a mixture of 16% paraformaldehyde/PBS. This mixture was incubated at 4 °C overnight.

On the following day, the pooled sample was washed and then 1× normalization “beta” beads were added and cells were strained into a new FACS tube. Cells were run on a Helios (Fluidigm) machine using the SuperSampler attachment. Following the run, the data were normalized to the normalization bead signal. Then, data was run through de-barcoder software (MATLAB; bead distance 0.2) and individual .FCS files were gated and analyzed manually using ImmuneAtlas and Cytobank platforms. tSNE plots were generated for filtered singlet/leukocyte (i.e., red blood cells excluded) populations for each file and formed into a composite plot. In brief, .FCS files were arcsinh transformed (a = 0, b = 1/5, c = 0) and 3000 randomly selected events were sampled for each file. tSNE was computed on the aggregated matrix with all events (3000 × 12 = 36,000). The following parameters were used for tSNE in the RTsne package (https://CRAN.R-project.org/package=Rtsne): perplexity = 30, max_iter = 500. Then, the transformed coordinate events were mapped back to the sample they come from, as well as composite maps for each condition and the total population.

### Statistical analysis

Hourly-based sleep comparisons were analyzed by a linear mixed-effects model followed by Šidák’s multiple comparisons as described in a similar counterbalanced crossover design (*55*). One-way ANOVA followed by Šidák’s multiple comparisons was used to analyze alteration of sleep/wake amount for different experimental paradigms, and latency to sleep onset. Paired student’s *t*-test was used for sleep/wake amount comparison between non-Stim and Stim datasets. CyTOF data was analyzed with Holm-Šidák test. In figures, *, **, ***, **** and ^†^ indicate *P* < 0.05, *P* < 0.01, *P* < 0.005, *P* < 0.001, and *P* < 0.0005 respectively. All values were reported as mean ± SEM.

## Acknowledgments

We thank de Lecea lab members for discussion, and A. Khan for excellent technical assistance. We also thank A. Olson and the Stanford Neuroscience Microscopy Service, NIH NS069375 for imaging technical assistance.

## Author contributions

S.-B.L. and L.d.L. conceived and designed the research. S.-B.L, J.C.B. and H.Y. performed the experiments and analyzed the data. J.H. assisted in CyTOF data analysis. B.G. supervised the CyTOF experiment and data analysis. L.d.L supervised the entire project. S.-B.L. wrote the manuscript with contributions from J.C.B, B.G. and L.d.L.

## Data and materials availability

All data needed to evaluate the conclusions in the paper are present in the paper and/or the Supplementary Materials. Additional data related to this paper may be requested with reasonable considerations.

---

**Fig. S1.**
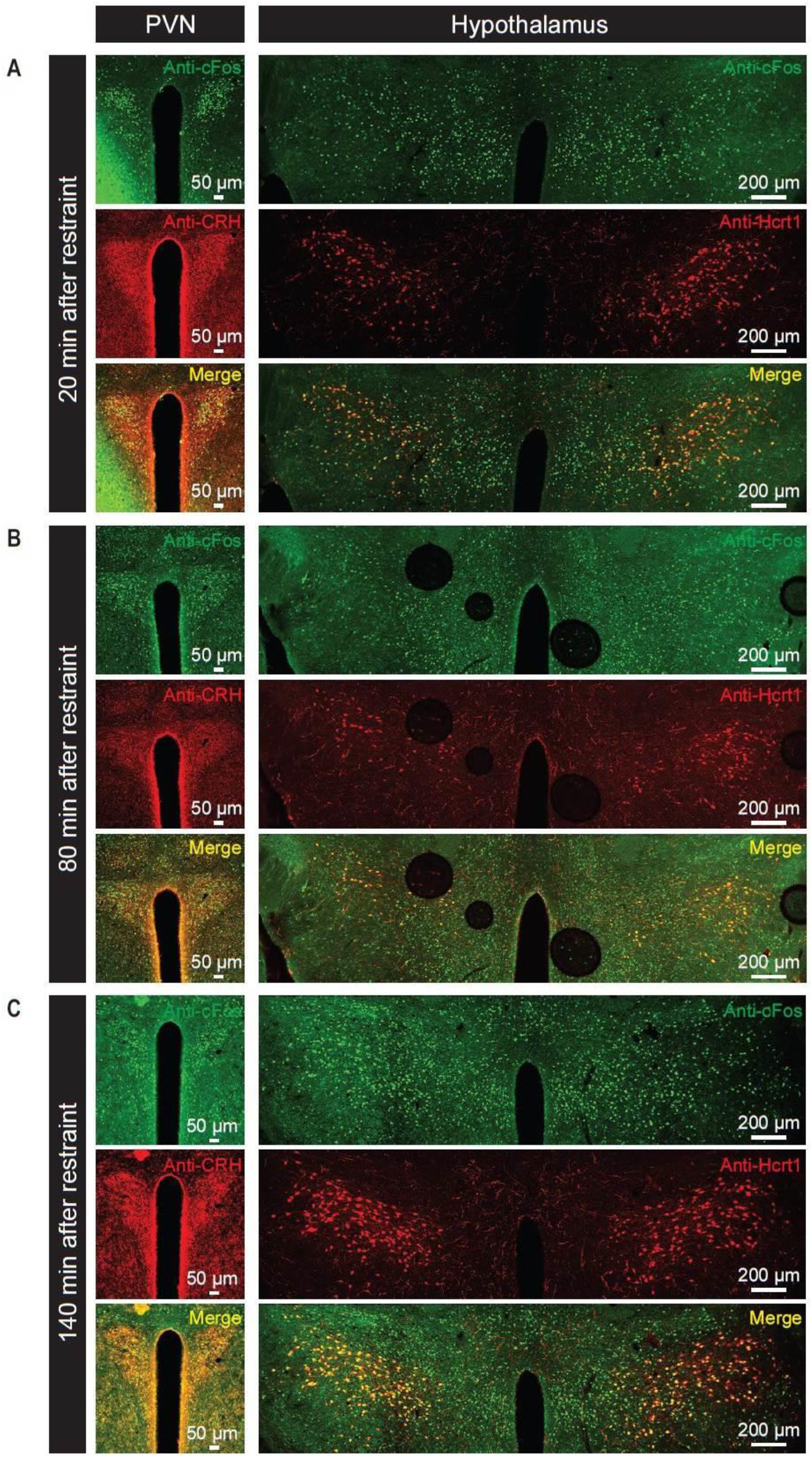
Restraint stress triggered cFos expression in CRH^**PVN**^ **and Hcrt**^**LH**^ **neurons**. A 10 min restraint stress session in animals’ home cages induced cFos expression in both CRH^PVN^ and Hcrt^LH^ neurons as validated at 20min (A), 80 min (B), and 140 min (C) after restraint onset (n = 3 each group).

**Fig. S2.**
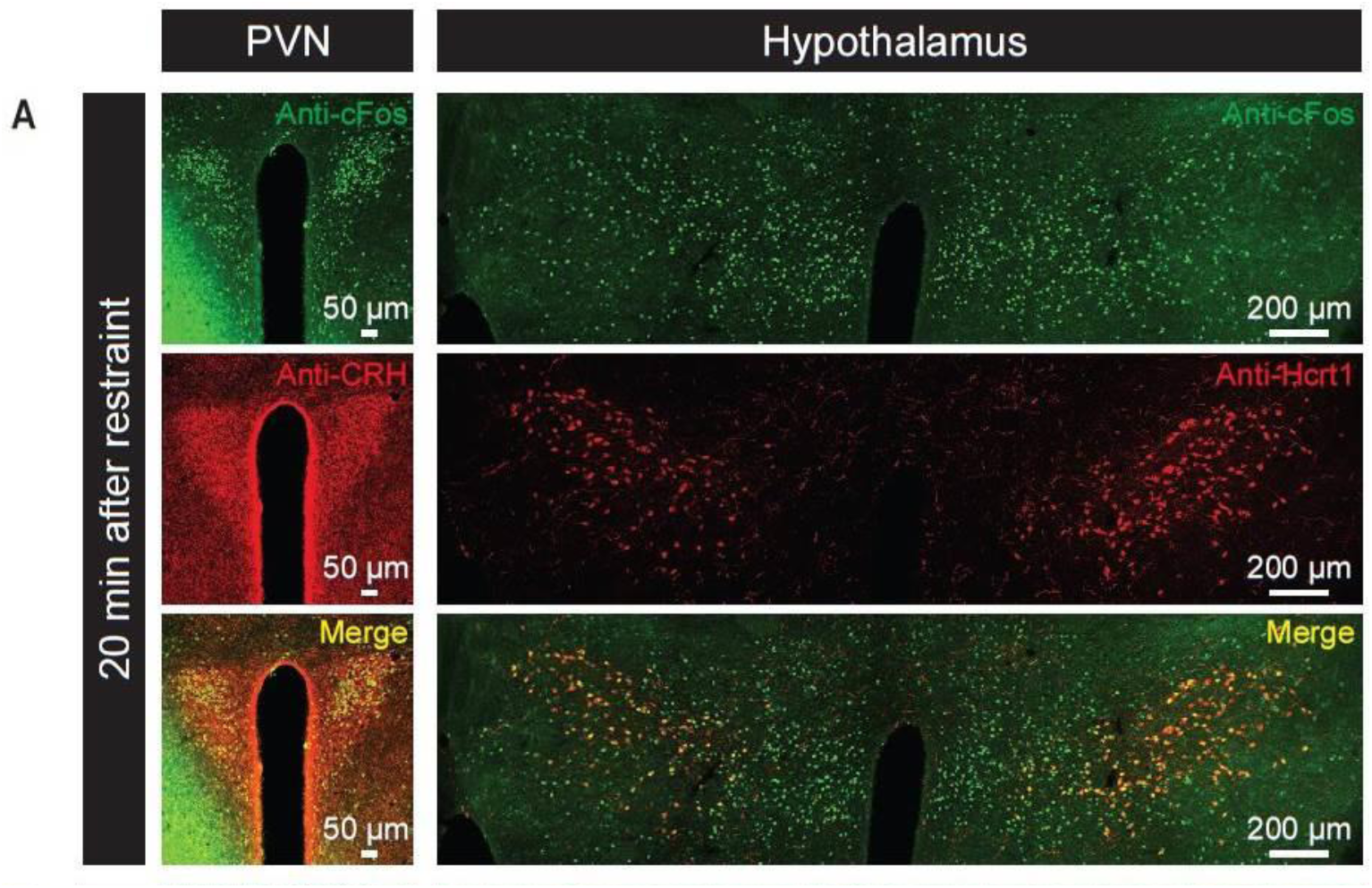
AAV-Retro vector validated projections from PVN to LH in CRH::Cre, AVP::Cre, and OXT::Cre mouse lines. (**A-E**) Representative slice containing PVN neurons labeled by AAV-Retro (**A**), antibody staining against CRH (**B**), merged slice (**C**), cell counts of AAV-Retro labeled neurons, CRH- positive neurons and both-positive neurons (**D**), and percentages of AAV-Retro labeled neurons positive for CRH staining and CRH neurons labeled by AAV-Retro (**E**) in CRH::Cre mice injected with AAV-Retro vectors in LH. (**F**-**J**) Representative slice containing PVN neurons labeled by AAV-Retro (**F**), antibody staining against AVP (**G**), merged slice (**H**), cell counts of AAV-Retro labeled neurons, AVP-positive neurons and both-positive neurons (**I**), and percentages of AAV-Retro labeled neurons positive for AVP staining and AVP neurons labeled by AAV-Retro (**J**) from AVP::Cre mice injected with AAV-Retro vectors in LH. (**K**-**O**) Representative slice containing PVN neurons labeled by AAV-Retro (**K**), antibody staining against OXT (**L**), merged slice (**M**), cell counts of AAV-Retro labeled neurons, OXT-positive neurons and both-positive neurons (**N**), and percentages of AAV-Retro labeled neurons positive for OXT staining and OXT neurons labeled by AAV-Retro (**O**) from OXT::Cre mice injected with AAV-Retro vectors to LH (n = 5 for each group).

**Fig. S3.**
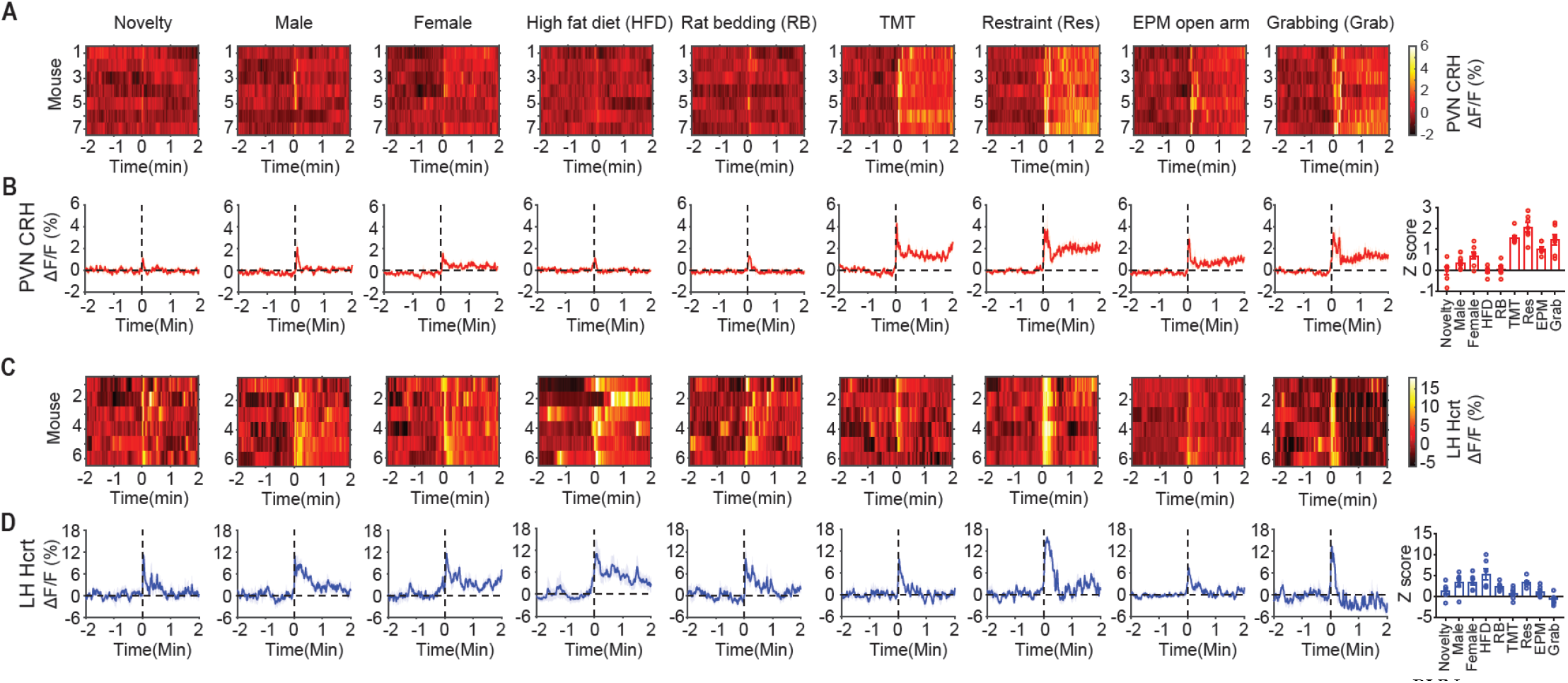
CRH^**PVN**^ **and Hcrt**^**LH**^ **neuronal activity during exposure to salient stimuli**. (**A**&**B**) CRH^PVN^ neurons and (**C**&**D**) Hcrt^LH^ neurons respond to distinct stimuli. Note aversive stimuli including predator fox odor (TMT), restraint stress (Res), elevated plus maze (EPM) open arm (mimicking anxiety condition) and grabbing (Grab) lead to strong CRH^PVN^ neuronal activities. Hcrt^LH^ neurons respond strongly to social, appetitive, and restraint stimuli (**A**&**B**: n = 7, **C**&**D**: n = 6).

**Fig. S4.**
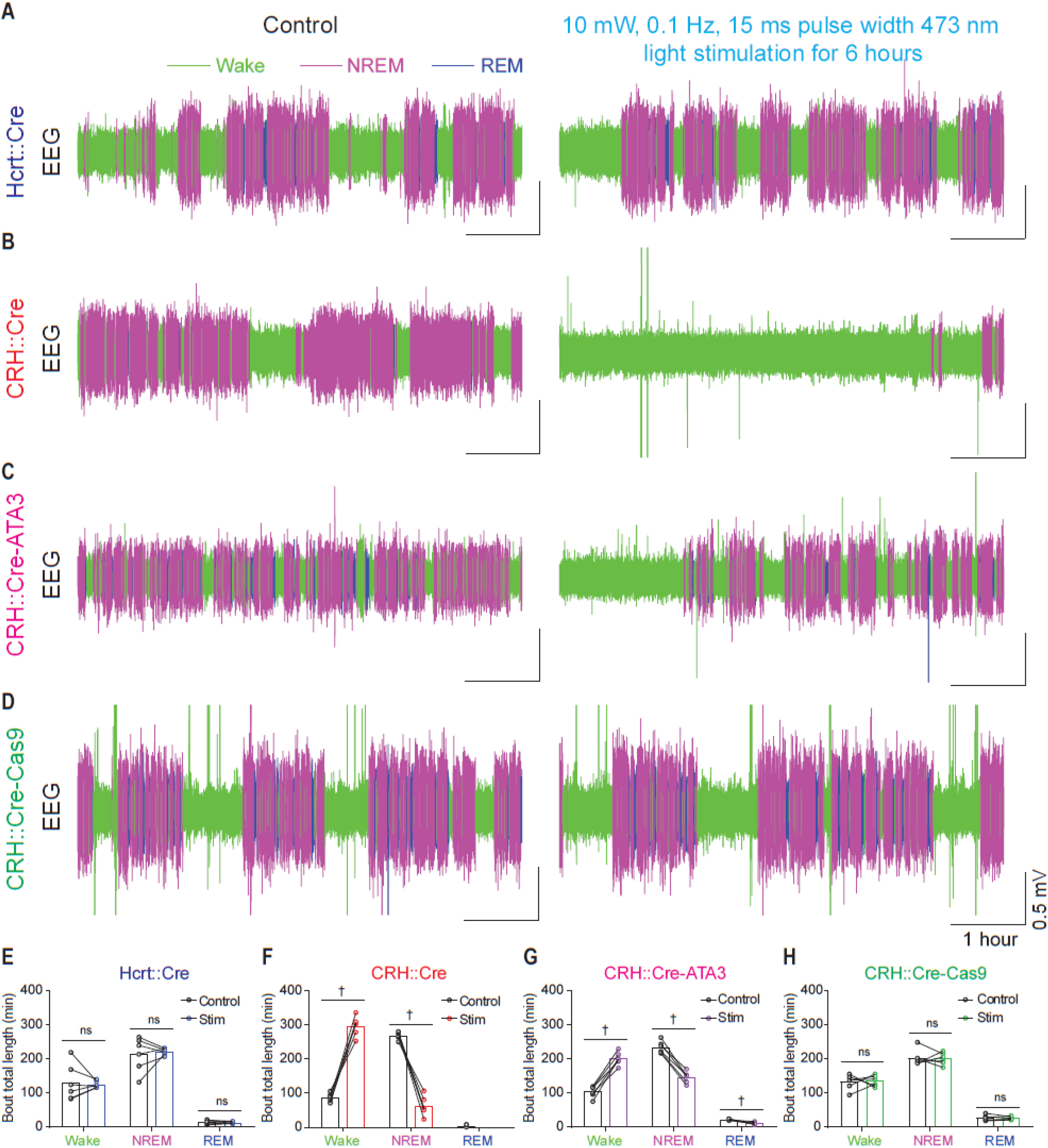
Prolonged mild optogenetic stimulation (10 mW, 0.**1 Hz, 15 ms 473 nm light pulse for 6 hours) of CRH**^**PVN**^ **neurons projecting to LH elicited hyperarousal/insomnia-like state**. (**A-D**) Representative EEG traces for control and optogenetic stimulation of Hcrt^LH^ neurons (**A**), LH-projecting CRH^PVN^ neurons in CRH::Cre mice (**B**), LH-projecting CRH^PVN^ neurons in CRH::Cre-ATA3 mice (**C**) and LH-projecting CRH^PVN^ neurons with CRH gene disrupted in CRH::Cre-Cas9 mice (**D**). (**E-H**), Comparison of total amount of wake, NREM sleep, REM sleep between control and optogenetic stimulation for Hcrt::Cre (**E**), CRH::Cre (**F**), CRH::Cre-ATA (**G**) and CRH::Cre-Cas9 (**H**) mice in the paradigm as described in **a** (paired *t*-test between Non-Stim and Stim, ^†^*P* < 0.0005, n = 6 mice for each group).

**Fig. S5.**
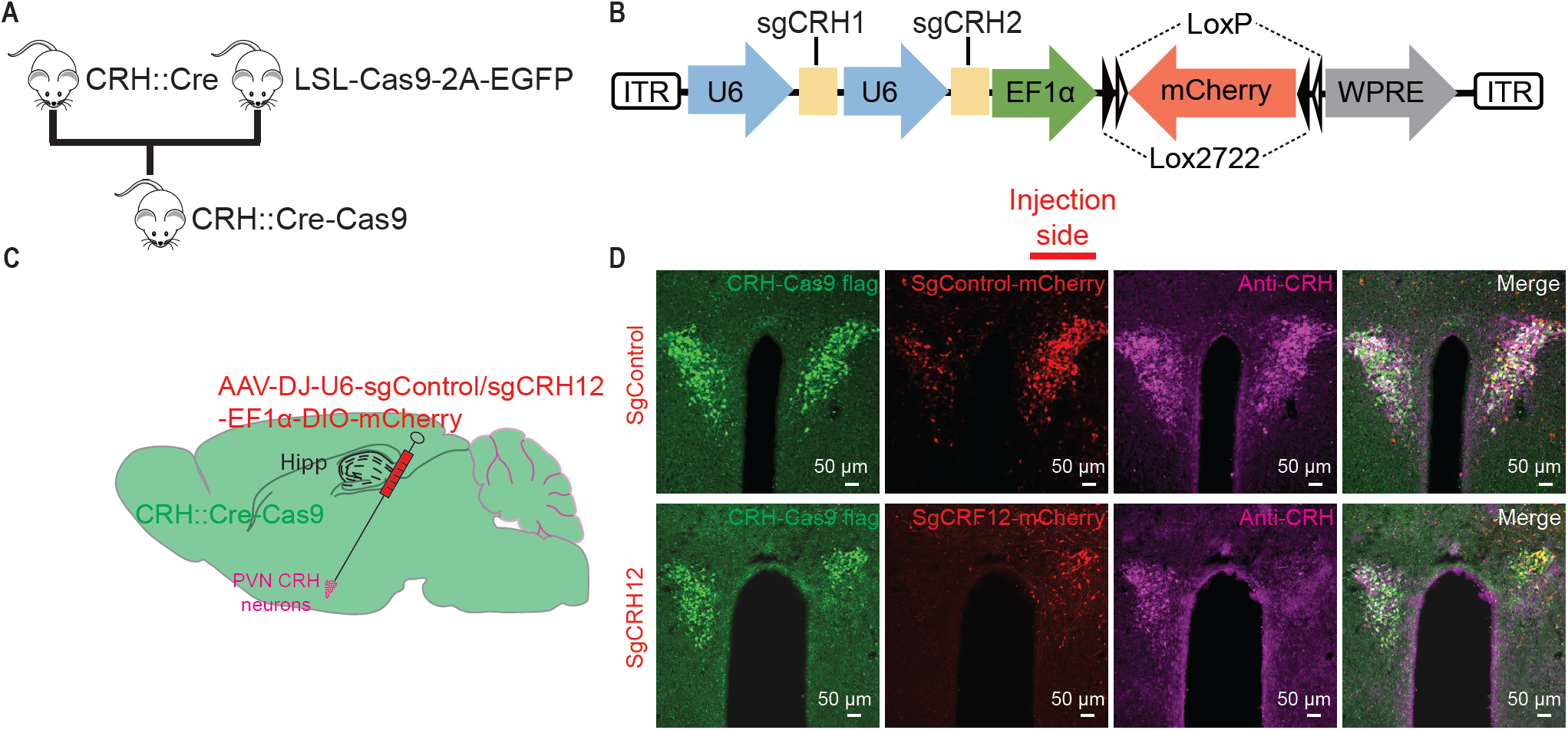
CRISPR-Cas9-mediated *crh* gene disruption in CRH^**PVN**^ **neurons of CRH::Cre-Cas9 mice.** **(A)** Generation of CRH::Cre-Cas9 mice. **(B)** Schematic of AAV sgCRH12 vector design. **(C)** Schematic of AAV-DJ-U6- sgControl/sgCRH12-EF1α-DIO-mCherry to unilateral PVN of CRH::Cre-Cas9 mice (n = 3, AP: −0.90 mm, ML: ±0.30 mm, DV: −4.5 + -4.8 mm, 0.3 μl each depth). **(D)** Representative slices from CRH::Cre-Cas9 mice injected with AAV vectors carrying sgControl or sgCRH12. Note the intact CRH staining on the contralateral side without viral infection in the sgCRH12 panel.

**Fig. S6.**
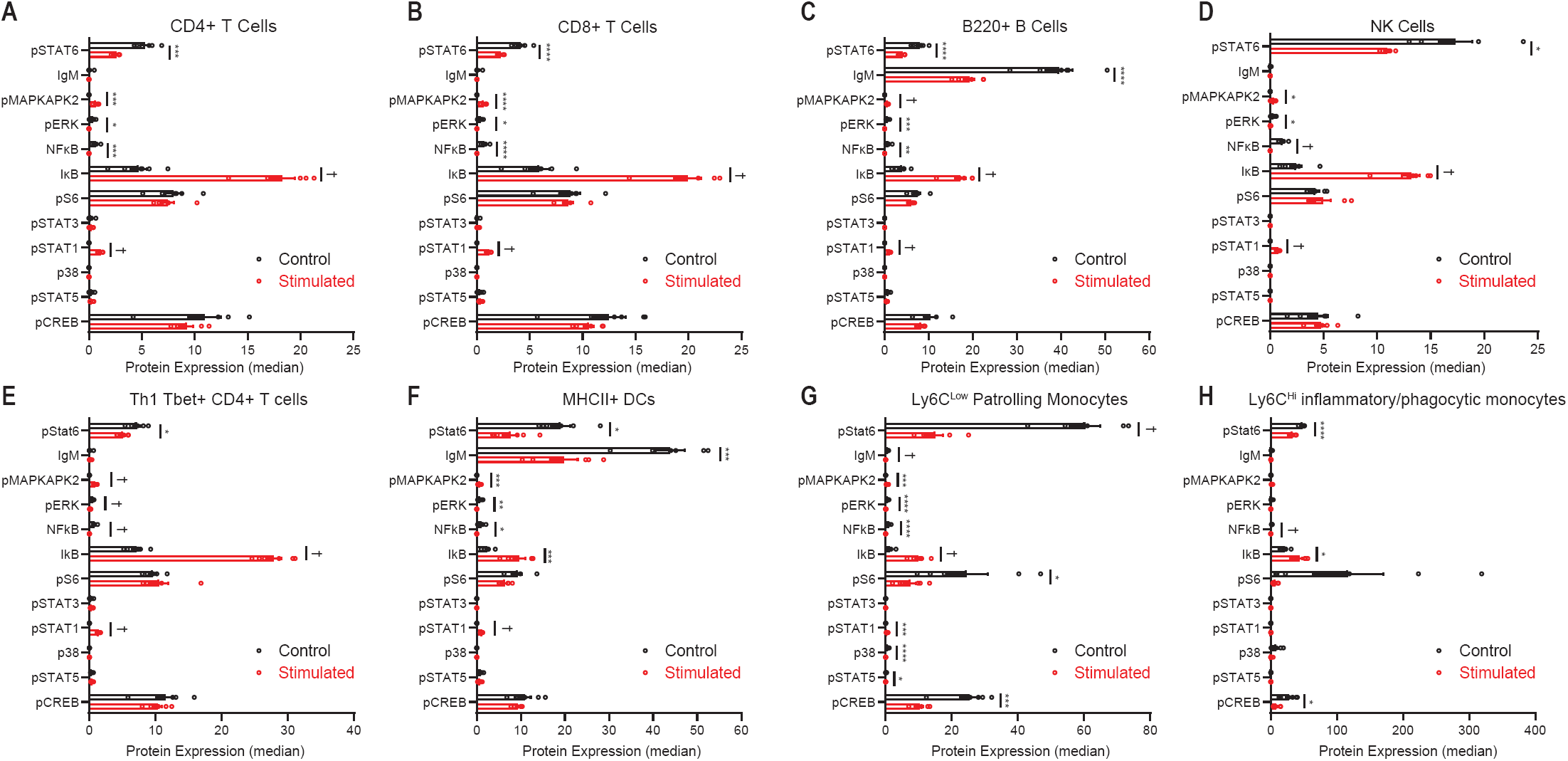
Optogenetic stimulation of CRH^**PVN**^ **neurons triggers dynamic changes in circulating immune cell intracellular signaling cascades.** **(A)** CD4+ T cells altered expression of pSTAT6 (control mean: 5.278; stimulation mean: 2.569; t ratio = 5.938, adjusted \*\*\**P* = 0.001148), pMAPKAPK2 (control mean: 0; stimulation mean: 0.655; t ratio = 6.048, adjusted \*\*\**P* = 0.001115), pERK (control mean: 0.3583, stimulation mean: 0, t ratio = 4.044, adjusted \**P* = 0.014003), NFκB (control mean: 0.6279, stimulation: 0; t ratio = 5.259, adjusted \*\*\**P* = 0.002578), IκB (control mean: 4.716; stimulation mean: 18.25; t ratio = 9.262, adjusted ^†^*P* = 0.000032), and pSTAT1 (control mean: 0; stimulation mean: 1.103, t ratio = 11.81, adjusted ^†^*P* = 0.000004). **(B)** CD8+ T cells had similar responses to optogenetic stimulation, pSTAT6 (control mean: 4.154; stimulation mean: 2.226; t ratio = 6.095, adjusted \*\*\*\**P* = 0.000996), pMAPKAPK2 (control mean: 0; stimulation mean: 0.6406, t ratio = 5.955, adjusted \*\*\*\**P* = 0.000996), pERK (control mean: 0.3365, stimulation mean: 0, t ratio = 4.148, adjusted \**P* = 0.011861), NFκB (control mean: 0.7248; stimulation mean: 0, t ratio = 6.133, adjusted \*\*\*\**P* = 0.000996), IκB (control mean: 5.893; stimulation mean: 19.89; t ratio = 8.77, adjusted ^†^*P* = 0.000052), and pSTAT1 (control mean: 0, stimulation mean: 1.142, t ratio = 13.23, adjusted ^†^*P* = 0.000001). **(C)** Additional changes in circulating B cells (B220+) were observed, including pSTAT6 (control mean: 7.963; stimulation mean: 4.026; t ratio = 5.83, adjusted \*\*\*\**P* = 0.000995), IgM (control mean: 39.55, stimulation mean: 19.18, t ratio = 6.308, adjusted \*\*\*\**P* = 0.000617), pMAPKAPK2 (control mean: 0, stimulation mean: 0.6665; t ratio = 8.724, adjusted ^†^*P* = 0.000044), pERK (control mean: 0.6719; stimulation mean: 0; t ratio = 5.610; adjusted \*\*\**P* = 0.001122), NFκB (control mean: 0.8489; stimulation mean: 0; t ratio = 4.257; adjusted \*\**P* **=** 0.006669), IκB (control mean: 3.894; stimulation mean: 17.19; t ratio = 9.626, adjusted ^†^*P* = 0.00002), and pSTAT1 (control mean: 0; stimulation mean: 1.172; t ratio = 13.11, adjusted ^†^*P* = 0.000001). **(D)** Additional changes were observed in circulating NK cells, pSTAT6 (control mean: 17.32; stimulation mean: 10.91; t ratio = 4.032, adjusted \**P* = 0.013537), pMAPKAPK2 (control mean: 0; stimulation mean: 0.3415; t ratio = 4.065, adjusted \**P* = 0.013537), pERK (control mean: 0.2802; stimulation mean: 0.01361; t ratio = 3.120, adjusted \**P* = 0.042791), NFκB (control mean: 1.149, stimulation mean: 0, t ratio = 8.253, adjusted ^†^*P* = 0.000063), IκB (control mean: 2.397, stimulation mean: 13.17, t ratio = 10.91, adjusted ^†^*P* = 0.00006), and pSTAT1 (control mean: 0, stimulation mean: 0.7538; t ratio = 14.2, adjusted ^†^*P* < 0.000001). **(E)** Alteration in Th1 Tbet+ CD4+ T cells including pSTAT6 (control mean: 7.177; stimulation mean: 5.069; t ratio = 3.526, adjusted \**P* = 0.032434), pMAPKAPK2 (control mean: 0; stimulation mean: 0.9113; t ratio = 8.570, adjusted ^†^*P* = 0.000058), pERK (control mean: 0.5380; stimulation mean: 0.1302; t ratio = 7.852, adjusted ^†^*P* = 0.000097), NFκB (control mean: 0.7856, stimulation mean: 0, t ratio = 8.152, adjusted ^†^*P* = 0.000080), IκB (control mean: 7.099, stimulation mean: 27.89, t ratio = 16.47, adjusted ^†^*P* < 0.000001), and pSTAT1 (control mean: 0, stimulation mean: 1.444; t ratio = 16.53, adjusted ^†^*P* < 0.000001). **(F)** Alteration in MHCII+ DCs including pSTAT6 (control mean: 19.02; stimulation mean: 7.716; t ratio = 4.049, adjusted \**P* = 0.011581), IgM (control mean: 43.87, stimulation mean: 19.82, t ratio = 5.298, adjusted \*\*\**P* = 0.002785), pMAPKAPK2 (control mean: 0, stimulation mean: 0.6783; t ratio = 5.387, adjusted \*\*\**P* = 0.002761), pERK (control mean: 0.7675; stimulation mean: 0; t ratio = 4.443; adjusted \*\**P* = 0.007468), NFκB (control mean: 1.032; stimulation mean: 0; t ratio = 3.741; adjusted \**P* = 0.015285), IκB (control mean: 2.434; stimulation mean: 9.693; t ratio = 5.130, adjusted \*\*\**P* = 0.003107), and pSTAT1 (control mean: 0; stimulation mean: 1.182; t ratio = 15.37, adjusted ^†^*P* < 0.000001). **(G)** Alteration in MHCII+ DCs including pSTAT6 (control mean: 60.33; stimulation mean: 14.94; t ratio = 8.598, adjusted ^†^*P* = 0.000062), IgM (control mean: 0.6686, stimulation mean: 0.01085, t ratio = 8.703, adjusted ^†^*P* = 0.000061), pMAPKAPK2 (control mean: 0, stimulation mean: 0.5179; t ratio = 4.272, adjusted \*\*\**P* = 0.004887), pERK (control mean: 0.6427; stimulation mean: 0.03657; t ratio = 6.487; adjusted \*\*\*\**P* = 0.000561), NFκB (control mean: 0.9809; stimulation mean: 0; t ratio = 6.301; adjusted \*\*\*\**P* = 0.000623), IκB (control mean: 1.250; stimulation mean: 9.956; t ratio = 7.986, adjusted ^†^*P* = 0.000108), pS6 (control mean: 24.60; stimulation mean: 7.506; t ratio = 2.643; adjusted \**P* = 0.048614), pSTAT1 (control mean: 0; stimulation mean: 0.4378; t ratio = 5.189, adjusted \*\*\**P* = 0.002036), p38 (control mean: 0.6725; stimulation mean: 0; t ratio = 6.278, adjusted \*\*\*\**P* = 0.000623), pSTAT5 (control mean: 0.1170; stimulation mean: 0.0009726; t ratio = 2.328, adjusted \**P* = 0.048614) and CREB (control mean: 25.39; stimulation mean: 10.40; t ratio = 5.023, adjusted \*\*\**P* = 0.002076). **(H)** pSTAT6 (control mean: 48.10; stimulation mean: 33.03; t ratio = 6.724, adjusted \*\*\*\**P* = 0.000572), NFκB (control mean: 1.651, stimulation mean: 0.03220, t ratio = 8.715, adjusted ^†^*P* = 0.000066), IκB (control mean: 21.25, stimulation mean: 43.71, t ratio = 4.394, adjusted \**P* = 0.013395), and CREB (control mean: 27.77, stimulation mean: 8.637; t ratio = 3.649, adjusted \**P* = 0.039519). (n = 6 mice/group; error bars represent SEM; Holm-Šidák Test; \**P* < 0.05, \*\**P* < 0.01, \*\*\**P* < 0.005, \*\*\*\**P* < 0.001, ^†^*P* < 0.0005).

**Table. S1.** Isotope conjugated antibodies used in CyTOF experiments.

| **Panel** | **Antigen** | **Atomic Symbol** | **Atomic Mass** |
| --- | --- | --- | --- |
| **Surface (Pre-MeOH)** | CD15 | Y | 89 |
|  | Ter-119 | In | 113 |
|  | CD45 | In | 115 |
|  | CD24 | La | 139 |
|  | CD27 | Ce | 140 |
|  | Ly6G | Pr | 141 |
|  | CD11c | Nd | 142 |
|  | CD11b | Nd | 143 |
|  | CD115 | Nd | 144 |
|  | CD4 | Nd | 145 |
|  | CD8a | Nd | 146 |
|  | PDCA1 | Sm | 147 |
|  | CD19 | Sm | 149 |
|  | CD3 | Sm | 152 |
|  | CD25 | Gd | 157 |
|  | CD16 | Gd | 158 |
| | TCR$\gamma\delta$ | Tb | 159 |
|  | CD62L | Gd | 160 |
|  | Ly6C | Dy | 162 |
|  | NK1.1 | Ho | 165 |
|  | IgM | Er | 169 |
|  | CD49b | Er | 170 |
|  | CD44 | Yb | 171 |
|  | CD38 | Yb | 173 |
|  | MHCII | Yb | 174 |
|  | CD127 | Lu | 175 |
|  | B220 | Lu | 176 |
| **Intracellular (Post-MeOH)** | pCREB | Nd | 148 |
|  | pSTAT5 | Nd | 150 |
|  | P38 | Eu | 151 |
|  | pSTAT1 | Eu | 153 |
|  | pSTAT3 | Sm | 154 |
|  | pS6 | Gd | 155 |
|  | Foxp3 | Gd | 156 |
|  | cPARP | Dy | 161 |
|  | Tbet | Dy | 163 |
| | IκB | Dy | 164 |
| | pNFκB | Er | 166 |
|  | pERK1/2 | Er | 167 |
|  | pMAPKAPK2 | Er | 168 |
|  | pSTAT6 | Yb | 172 |

## Reference

1. T. L. Sterley et al., Social transmission and buffering of synaptic changes after stress. Nat Neurosci 21, 393–403 (2018).

2. G. Cano, T. Mochizuki, C. B. Saper, Neural circuitry of stress-induced insomnia in rats. The Journal of neuroscience : the official journal of the Society for Neuroscience 28, 10167–10184 (2008).

3. J. N. Morey, I. A. Boggero, A. B. Scott, S. C. Segerstrom, Current Directions in Stress and Human Immune Function. Curr Opin Psychol 5, 13–17 (2015).

4. A. N. Vgontzas et al., Chronic insomnia is associated with nyctohemeral activation of the hypothalamic-pituitary-adrenal axis: clinical implications. J Clin Endocrinol Metab 86, 3787–3794 (2001).

5. A. N. Vgontzas et al., Chronic insomnia and activity of the stress system: a preliminary study. J Psychosom Res 45, 21–31 (1998).

6. A. R. Adamantidis, F. Zhang, A. M. Aravanis, K. Deisseroth, L. de Lecea, Neural substrates of awakening probed with optogenetic control of hypocretin neurons. Nature 450, 420–424 (2007).

7. M. E. Carter et al., Mechanism for Hypocretin-mediated sleep-to-wake transitions. Proc Natl Acad Sci U S A 109, E2635–2644 (2012).

8. M. E. Carter et al., Tuning arousal with optogenetic modulation of locus coeruleus neurons. Nat Neurosci 13, 1526–1533 (2010).

9. A. Eban-Rothschild, G. Rothschild, W. J. Giardino, J. R. Jones, L. de Lecea, VTA dopaminergic neurons regulate ethologically relevant sleep-wake behaviors. Nat Neurosci 19, 1356–1366 (2016).

10. J. R. Cho et al., Dorsal Raphe Dopamine Neurons Modulate Arousal and Promote Wakefulness by Salient Stimuli. Neuron 94, 1205–1219 e1208 (2017).

11. M. Xu et al., Basal forebrain circuit for sleep-wake control. Nat Neurosci 18, 1641–1647 (2015).

12. S. O. Irmak, L. de Lecea, Basal forebrain cholinergic modulation of sleep transitions. Sleep 37, 1941–1951 (2014).

13. Y. Yuan et al., Reward Inhibits Paraventricular CRH Neurons to Relieve Stress. Curr Biol 29, 1243–1251 e1244 (2019).

14. W. J. Giardino et al., Parallel circuits from the bed nuclei of stria terminalis to the lateral hypothalamus drive opposing emotional states. Nat Neurosci 21, 1084–1095 (2018).

15. K. H. Schulz, S. Gold, [Psychological stress, immune function and disease development. The psychoneuroimmunologic perspective]. Bundesgesundheitsblatt Gesundheitsforschung Gesundheitsschutz 49, 759–772 (2006).

16. S. C. Segerstrom, G. E. Miller, Psychological stress and the human immune system: a meta-analytic study of 30 years of inquiry. Psychol Bull 130, 601–630 (2004).

17. J. K. Kiecolt-Glaser, J. R. Dura, C. E. Speicher, O. J. Trask, R. Glaser, Spousal caregivers of dementia victims: longitudinal changes in immunity and health. Psychosom Med 53, 345–362 (1991).

18. K. Mizobe et al., Restraint stress-induced elevation of endogenous glucocorticoid suppresses migration of granulocytes and macrophages to an inflammatory locus. J Neuroimmunol 73, 81–89 (1997).

19. N. Tarcic, H. Ovadia, D. W. Weiss, J. Weidenfeld, Restraint stress-induced thymic involution and cell apoptosis are dependent on endogenous glucocorticoids. J Neuroimmunol 82, 40–46 (1998).

20. D. W. Cain, J. A. Cidlowski, Immune regulation by glucocorticoids. Nat Rev Immunol 17, 233–247 (2017).

21. J. Minkel et al., Sleep deprivation potentiates HPA axis stress reactivity in healthy adults. Health Psychol 33, 1430–1434 (2014).

22. A. Zimprich et al., A robust and reliable non-invasive test for stress responsivity in mice. Frontiers in behavioral neuroscience 8, (2014).

23. A. Ramot et al., Hypothalamic CRFR1 is essential for HPA axis regulation following chronic stress. Nat Neurosci 20, 385–388 (2017).

24. Y. C. Saito et al., Monoamines Inhibit GABAergic Neurons in Ventrolateral Preoptic Area That Make Direct Synaptic Connections to Hypothalamic Arousal Neurons. Journal of Neuroscience 38, 6366–6378 (2018).

25. E. M. Callaway, L. Luo, Monosynaptic Circuit Tracing with Glycoprotein-Deleted Rabies Viruses. The Journal of neuroscience : the official journal of the Society for Neuroscience 35, 8979–8985 (2015).

26. N. R. Wall, I. R. Wickersham, A. Cetin, M. De La Parra, E. M. Callaway, Monosynaptic circuit tracing in vivo through Cre-dependent targeting and complementation of modified rabies virus. P Natl Acad Sci USA 107, 21848–21853 (2010).

27. D. G. Tervo et al., A Designer AAV Variant Permits Efficient Retrograde Access to Projection Neurons. Neuron 92, 372–382 (2016).

28. H. S. Knobloch et al., Evoked axonal oxytocin release in the central amygdala attenuates fear response. Neuron 73, 553–566 (2012).

29. M. G. Lee, O. K. Hassani, B. E. Jones, Discharge of identified orexin/hypocretin neurons across the sleep-waking cycle. The Journal of neuroscience : the official journal of the Society for Neuroscience 25, 6716–6720 (2005).

30. B. Y. Mileykovskiy, L. I. Kiyashchenko, J. M. Siegel, Behavioral correlates of activity in identified hypocretin/orexin neurons. Neuron 46, 787–798 (2005).

31. J. Kim et al., Rapid, biphasic CRF neuronal responses encode positive and negative valence. Nat Neurosci, (2019).

32. J. S. Kim, S. Y. Han, K. J. Iremonger, Stress experience and hormone feedback tune distinct components of hypothalamic CRH neuron activity. Nature communications 10, 5696 (2019).

33. J. Hara et al., Genetic ablation of orexin neurons in mice results in narcolepsy, hypophagia, and obesity. Neuron 30, 345–354 (2001).

34. H. Yamaguchi, F. W. Hopf, S. B. Li, L. de Lecea, In vivo cell type-specific CRISPR knockdown of dopamine beta hydroxylase reduces locus coeruleus evoked wakefulness. Nature communications 9, 5211 (2018).

35. C. P. Romanowski et al., Central deficiency of corticotropin-releasing hormone receptor type 1 (CRH-R1) abolishes effects of CRH on NREM but not on REM sleep in mice. Sleep 33, 427–436 (2010).

36. R. Winsky-Sommerer et al., Interaction between the corticotropin-releasing factor system and hypocretins (Orexins): A novel circuit mediating stress response. Journal of Neuroscience 24, 11439–11448 (2004).

37. S. C. Bendall et al., Single-cell mass cytometry of differential immune and drug responses across a human hematopoietic continuum. Science 332, 687–696 (2011).

38. R. Glaser, J. K. Kiecolt-Glaser, Stress-induced immune dysfunction: implications for health. Nat Rev Immunol 5, 243–251 (2005).

39. N. Auphan, J. A. DiDonato, C. Rosette, A. Helmberg, M. Karin, Immunosuppression by glucocorticoids: inhibition of NF-kappa B activity through induction of I kappa B synthesis. Science 270, 286–290 (1995).

40. R. I. Scheinman, P. C. Cogswell, A. K. Lofquist, A. S. Baldwin, Jr., Role of transcriptional activation of I kappa B alpha in mediation of immunosuppression by glucocorticoids. Science 270, 283–286 (1995).

41. A. Biola et al., The glucocorticoid receptor and STAT6 physically and functionally interact in T-lymphocytes. FEBS letters 487, 229–233 (2000).

42. A. E. Coutinho, K. E. Chapman, The anti-inflammatory and immunosuppressive effects of glucocorticoids, recent developments and mechanistic insights. Mol Cell Endocrinol 335, 2–13 (2011).

43. M. Wang et al., Acute restraint stress enhances hippocampal endocannabinoid function via glucocorticoid receptor activation. Journal of psychopharmacology 26, 56–70 (2012).

44. C. J. Cook, Stress induces CRF release in the paraventricular nucleus, and both CRF and GABA release in the amygdala. Physiol Behav 82, 751–762 (2004).

45. E. Elliott, G. Ezra-Nevo, L. Regev, A. Neufeld-Cohen, A. Chen, Resilience to social stress coincides with functional DNA methylation of the Crf gene in adult mice. Nat Neurosci 13, 1351–1353 (2010).

46. J. C. Zant et al., Cholinergic Neurons in the Basal Forebrain Promote Wakefulness by Actions on Neighboring Non-Cholinergic Neurons: An Opto-Dialysis Study. The Journal of neuroscience : the official journal of the Society for Neuroscience 36, 2057–2067 (2016).

47. D. J. Taylor, K. L. Lichstein, H. H. Durrence, Insomnia as a health risk factor. Behav Sleep Med 1, 227–247 (2003).

48. L. A. Wang, D. H. Nguyen, S. W. Mifflin, Corticotropin-releasing hormone projections from the paraventricular nucleus of the hypothalamus to the nucleus of the solitary tract increase blood pressure. J Neurophysiol 121, 602–608 (2019).

49. E. M. Friedman, M. R. Irwin, A role for CRH and the sympathetic nervous system in stress-induced immunosuppression. Annals of the New York Academy of Sciences 771, 396–418 (1995).

50. M. L. Hanke, N. D. Powell, L. M. Stiner, M. T. Bailey, J. F. Sheridan, Beta adrenergic blockade decreases the immunomodulatory effects of social disruption stress. Brain Behav Immun 26, 1150–1159 (2012).

51. L. de Lecea et al., The hypocretins: hypothalamus-specific peptides with neuroexcitatory activity. Proc Natl Acad Sci U S A 95, 322–327 (1998).

52. T. Sakurai et al., Orexins and orexin receptors: a family of hypothalamic neuropeptides and G protein-coupled receptors that regulate feeding behavior. Cell 92, 573–585 (1998).

53. K. Labun, T. G. Montague, J. A. Gagnon, S. B. Thyme, E. Valen, CHOPCHOP v2: a web tool for the next generation of CRISPR genome engineering. Nucleic Acids Res 44, W272–276 (2016).

54. F. A. Ran et al., Genome engineering using the CRISPR-Cas9 system. Nat Protoc 8, 2281–2308 (2013).

55. S. B. Li, N. Nevárez, W. J. Giardino, L. de Lecea, Optical probing of orexin/hypocretin receptor antagonists. Sleep 41, (2018).

